# Regulation of BCR-mediated Ca^2+^ mobilization by MIZ1-TIMBIM4 safeguards IgG1^+^ GC B cell positive selection

**DOI:** 10.1101/2023.07.18.549490

**Authors:** Lingling Zhang, Amparo Toboso-Navasa, Arief Gunawan, Abdouramane Camara, Rinako Nakagawa, Katja Finsterbusch, Probir Chakravarty, Rebecca Newman, Yang Zhang, Martin Eilers, Wack Andreas, Pavel Tolar, Kai-Michael Toellner, Dinis Pedro Calado

## Abstract

The transition from IgM to affinity-matured IgG antibodies is vital for effective humoral immunity. This is facilitated by germinal centers (GCs) through affinity maturation and preferential accumulation of IgG^+^ B cells over IgM^+^ B cells. However, it is not known whether the positive selection of the different immunoglobulin isotypes within GCs varies in its dependency on specific transcriptional mechanisms. Here, we identified IgG1^+^ GC B cell transcription factor dependency using CRISPR-Cas9 and conditional mouse genetics. We found that MIZ1 was specifically required for IgG1^+^ GC B cell survival during positive selection, whereas IgM^+^ GC B cells were largely independent. Mechanistically, MIZ1 induced TMBIM4, an ancestral anti-apoptotic protein that regulated inositol trisphosphate receptor mediated Ca^2+^ mobilization downstream of IgG1. The MIZ1-TMBIM4 axis prevented mitochondrial dysfunction-induced IgG1^+^ GC cell death caused by excessive Ca^2+^ accumulation. This study uncovers a unique immunoglobulin isotype-specific dependency, on a hitherto unidentified mechanism in GC positive selection.

## Introduction

For an effective humoral immune response to occur, a transition is required from IgM to affinity-matured IgG antibodies^1–3^. This transition is achieved through the accumulation of IgG^+^ B cells during germinal center (GC) reactions and by the enhancement of antibody affinity towards the antigen^4,5^. The importance of this process is underscored by the occurrence of recurrent infections and unfavorable clinical outcomes observed in patients with hyper-IgM syndrome. These individuals carry e.g., mutations in activation-induced cytidine deaminase (AID)^6–8^, which renders B cells deficient in class switch recombination (CSR) and somatic hypermutation (SHM).

GCs are central for affinity maturation, leading to the generation of plasma cells (PCs) that produce antibodies with enhanced affinity, and memory B cells (MBCs) that rapidly differentiate into PCs upon antigen re-challenge^9–12^. These specialized structures form in secondary lymphoid organs during immune responses that are antigen and T cell dependent^13,14^. GCs are comprised of two distinct regions known as the light zone (LZ) and dark zone (DZ). Somatic hypermutation, driven by AID, takes place in the DZ. In the LZ, GC B cells undergo a selection process that is partly based on the affinity of their B cell receptors (BCRs) for antigen immune complexes presented by follicular dendritic cells (FDCs)^13,14^. Positively selected GC B cells (∼5-10% of the LZ) represent those that have productively extracted antigen from FDCs via their BCR and have presented it as peptides on major histocompatibility complex (pMHC) eliciting T follicular helper cell (TFH)-help^13,15–17^. Our research, and other studies, identified the expression of MYC as a marker of positively selected GC B cells^17–19^.

In GCs there is also a preferential accruement of IgG^+^ over IgM^+^ B cells as the GC reaction progresses^20,21^. This cannot be attributed to IgM to IgG CSR because it predominantly occurs before B cells enter a GC reaction and is infrequent in established GCs^21–23^. Moreover, recent studies uncovered that the selection of B cells in GCs is a dynamic and multi-layered biological process that extends beyond BCR affinity-dependence *per se*^24–28^. As an example, parallel forms of positive selection in GC B cells, built on BCR affinity and the capacity of the immunoglobulin constant region to extract antigen from FDCs, seem required to ensure IgG1^+^ GC B cell positive selection^21^.

Still, the activity of GC B cell BCR signaling remains particularly enigmatic. Studies utilizing transgenic systems revealed that IgG BCRs exhibit more robust and sustained Ca^2+^ mobilization compared to IgM BCRs, which could be considered an advantage^29–33^. However, in GCs, BCR signaling is attenuated and rewired, resulting in compromised BCR-mediated Ca^2+^ mobilization^14,17,34,35^. Notably, there is currently an apparent contradiction between these findings and those of others using a transcriptional reporter of BCR signaling (i.e., Nur77 reporter)^36,37^, and advanced imaging techniques^38,39^. While these studies have not provided immunoglobulin isotype resolution, they raise intriguing questions regarding the activity of BCR-mediated Ca^2+^ mobilization in GC B cells.

Despite its significance, it is not known whether the positive selection of different immunoglobulin isotypes within GCs depends variably on specific transcriptional mechanisms. Here, we found that the transcription factor MIZ1 played an essential and specific role in safeguarding IgG1^+^ GC B cells during positive selection, whereas IgM^+^ GC B cells were largely independent. Mechanistically, MIZ1 activated the expression of TMBIM4, a newly identified target gene, member of an ancestral protein family with anti-apoptotic properties^40,41^. TMBIM4 effectively regulated Ca^2+^ release from the inositol trisphosphate receptor (IP3R)^42–45^ downstream of IgG1 BCR. Lack of the MIZ1-TMBIM4 axis, led to exacerbated Ca^2+^ mobilization specifically in IgG1^+^ GC B cells, resulting in mitochondrial dysfunction and cell death. Restoration of TMBIM4 expression *in vivo* rescued positive selection of MIZ1-deficient IgG1^+^ GC B cells. This study identifies for the first time a critical role for MIZ in regulating Ca^2+^ release and establishes the *in vivo* significance of the TMBIM4 anti-apoptotic factor. Significantly, this research provides evidence for a unique immunoglobulin isotype-specific dependency, on a previously unidentified transcriptional mechanism in GC positive selection.

## Results

### MIZ1 identified in a IgG1^+^ B cell CRISPR/Cas9 screen

We analyzed a CRISPR/Cas9 screen using *in vitro* derived GC B cells (iGB) to identify candidate transcription factors (TFs) required for the survival and/or proliferation of IgG1^+^ GC B cells during selection^46^. This genome-wide CRISPR screening was performed by constructing the sgRNA Cherry-Brie library and infecting Cas9-expressing naïve mouse B cells^46,47^. Briefly, IgM^+^ naïve mouse B cells were transduced and cultured on 40LB feeder cells with IL-4 to induce efficient CSR and generation of IgG1^+^ iGB cells. Antigen stimulation was then provided by adding anti-Igκ F(ab’)2, mimicking the signal required for B cell selection in GCs^48,17^. IgG1^+^ iGB cells were purified using flow cytometry and deep sequencing of sgRNAs was performed to identify depleted sgRNAs and assign CRISPR scores to genes (**Figure 1A**).

**Figure. 1.**
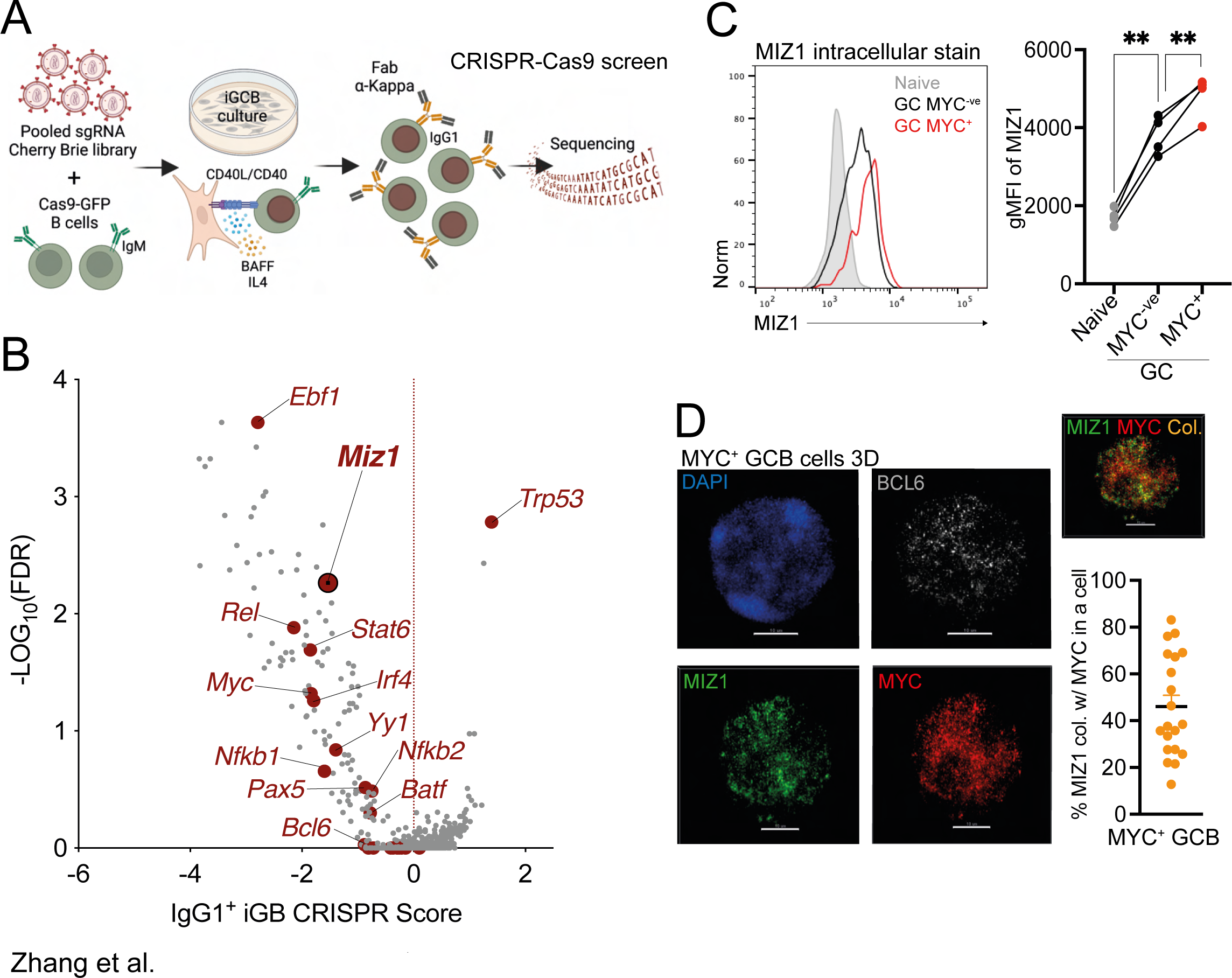
CRISPR screens identifies *Miz1* as an essential TF in IgG1^+^ B cells. **(A)** Schematic of CRISPR screen workflow and IgG1^+^ analysis strategy. **(B)** Volcano plot of TF gene CRISPR scores versus statistical significance corrected for false discovery rate (FDR) for all TFs (gray) and essential TFs (dark red). (C-D) C57BL/6 mice were i.v. immunized with SRBCs, 8-9 days after immunization spleens were collected. **(C)** Left, representative plot for MIZ1 expression levels in naive B cells, MYC^-ve^ and MYC^+^ GC B cells by intracellular stain. Right, representative data for MIZ1 expression levels in these cell populations, measured by gMFI. Cell populations from the same mouse were lined up, **, P ≤ 0.01 (multiple T test). **(D)** Immunofluorescence stain of freshly isolated GC B cells. GC B cells were negatively isolated from splenocytes, seeded onto chamber slides and fixed before further staining. Left, representative counterstain of DAPI (blue), anti-BCL6 antibody (grey), antiIZ1 antibody (green) and anti-MYC antibody (red) in one GC B cell, scale bars = 20 μM. Right, distribution of MIZ1 (green) and MYC (red) in the nucleus, colocalization of MIZ1 and MYC is highlighted as orange = Colocalization. Graph represents the percentage of MIZ1 co-localized with MYC in a single MYC^+^MIZ1^+^ GC B cell. Each dot represents one cell (n=20), small horizontal line represents mean and SEM.

Among the TFs identified in the CRISPR screen, *Ebf1* ranked the highest (**Figure 1B**). This finding was consistent with the established role of EBF1 in the overall maintenance of GC B cells and therefore not limited to IgG1^+^ GC B cells specifically^49^. Several identified TFs, including MYC, BCL6, PAX5, REL, BATF, YY1, and STAT6, also had similar characteristics^18,19,50–60^. While NFKB1, NFKB2, and IRF4 are not crucial for GC B cell maintenance, evidence suggests their depletion could reflect an impact on CSR^61–66^.

Among the genes identified in the CRISPR screen as crucial for IgG1^+^ B cell proliferation or maintenance was the TF MIZ1. We previously found high MIZ1 expression in MYC^+^ GC B cells^67^, which represent positively selected GC B cells^18,19^ (**Figure 1C**). Both MYC and MIZ1 function as transcriptional activators; however, they can also form complexes that repress the expression of MIZ1 target genes^67–70^. However, mice with disrupted MYC-MIZ1 complex formation showed no significant alterations in IgG1^+^ B cell positive selection within GCs^67^. This observation suggests that MIZ1 has roles in GC B cell selection, beyond its interaction with MYC. Notably, 3D imaging of freshly isolated positively selected GC B cells revealed that about half of MIZ1 molecules did not co-localize with MYC (**Figure 1D**, **S1A** and **Supplemental Video 1**). This indicates that a significant proportion of MIZ1 molecules operate independently of MYC, potentially facilitating MIZ1-dependent transcriptional activation.

### MIZ1 deficiency leads to loss of IgG1^+^ GC B cells *in vivo*

To study the role of MIZ1 in positively selected GC B cells, we utilized *Cγ1-cre*^71^ and a conditional allele with loxP sites flanking the MIZ1 POZ domain^72^, that is essential for transcriptional activity (hereafter called MIZ1^KO^). In addition, we used a *Rosa26* cre-loxP fluorescent reporter^73^ (*R26eYFP^stopFL^*) to complement the MIZ1^KO^ genetic system. This reporter enabled the identification of eYFP^+^ cells as an indicator of successful cre-mediated recombination. Mice carrying the *Cγ1-cre* and *R26eYFP^stopFL^* alleles were used as controls and are hereafter referred to as Ctrl (**Figure 2A**). Furthermore, we generated compound mutant mice that also harbored the SWHEL system, where B cells express a transgenic BCR specific to hen egg lysozyme (HEL)^74^.

**Figure. 2.**
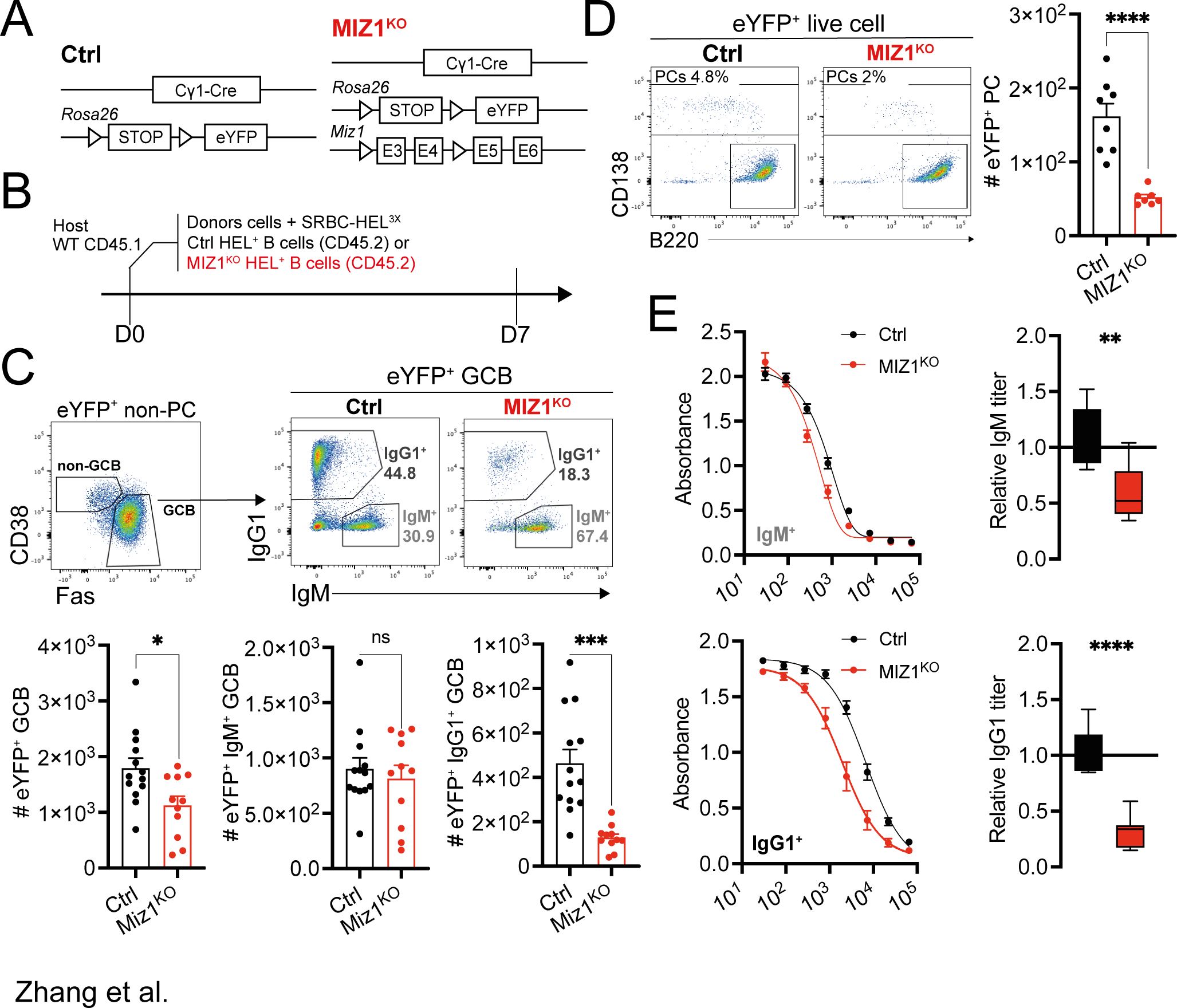
MIZ1 deficiency leads to the loss of IgG1^+^ GC B cell *in vivo*. **(A)** Schematics of allele combination in control (Ctrl) and experimental mice (MIZ1^KO^). An allele containing *loxP-stop-loxP-eYFP* (*loxP=* triangle) in the *Rosa26* locus is present in both Ctrl and MIZ1^KO^ mice. MIZ1^KO^ contains an *Miz1* allele with two *loxP* (triangle) flanking exon3 and exon4 that encode the BTB/POZ DNA binding domain. **(B)** SWHEL experiment schematics. **(C)** Top, gating strategy for IgG1^+^ and IgM^+^ GC B cell subsets generated from donor-derived reporter positive B cells. Bottom, graphs displaying cumulative data of relative cell numbers (numbers per 10^6^ splenocytes) for total GC B cells, and IgG1^+^ and IgM^+^ GC B cell subsets in, Ctrl (control, black); and MIZ1^KO^ (red). **(D)** Left, gating strategy for eYFP^+^ plasma cells (PCs). Right, graphs displaying cumulative data of relative cell numbers (numbers per 10^6^ splenocytes) for total PCs. **(E)** Left, ELISA for HEL-binding IgG1 and IgM antibodies in recipient mice. Right, fold changes of relative antibody titer of anti-HEL IgG1 (OD = 1), and anti-HEL IgM (OD = 0.4). Each symbol (C: Ctrl n = 12, MIZ1^KO^ n = 12; D: Ctrl n = 8, MIZ1^KO^ n = 7; E: Ctrl n = 7, MIZ1^KO^ n = 7) represents an individual mouse; small horizontal lines show mean and SEM. Data in (C-D) is representative of three to five experiments. Data in (E) is from two independent experiments. *, P ≤ 0.05; **, P ≤ 0.01; ***, P ≤ 0.001; ****, P ≤ 0.0001 (unpaired two-tailed Student’s t test. ns, not significant.)

First, we assessed the impact of POZ domain deletion on the functionality of MIZ1. Previous research demonstrated that truncated MIZ1 lacking the POZ domain is incapable of binding to chromatin or to other TFs^72^. Consistently, using 3D imaging on freshly isolated positively selected GC B cells, we observed a near complete loss in the co-localization of MYC and MIZ1 in MIZ1^KO^ compared to Ctrl (**Figure S1**). We next examined the impact of the loss of MIZ1’s transcriptional activity on GC B cell dynamics. Following established protocols^75^, we transferred CD45.2^+^ HEL-specific B cells from donor MIZ1^KO^ and Ctrl mice into CD45.1^+^ recipient mice. Subsequently, the recipient mice were immunized with HEL^3X^ antigen (**Figure 2B**). At day 7 post-immunization, flow cytometry analysis was performed on spleens. We focused on eYFP^+^ cells to identify GC B cells and quantified the populations of IgM^+^ and IgG1^+^ B cells (**Figure 2C**). Inactivation of MIZ1’s transcriptional activity resulted in a decrease in the total number of GC B cells, which is consistent with previous findings^76^. However, the number of IgM^+^ GC B cells remained unaffected. In contrast, MIZ1^KO^ mice exhibited a significant reduction in the number of IgG1^+^ GC B cells, with less than 25% of cells remaining compared to the Ctrl (**Figure 2C**). We further confirmed this observation in MIZ1^KO^ mice with a polyclonal BCR repertoire following SRBCs immunization (**Figure S2A**). These findings indicate that MIZ1 induced transcriptional activation is specifically required in IgG1^+^ B cells.

### MIZ1 is required for IgG1^+^ B cell accruement in GCs

The decrease in IgG1^+^ B cell numbers in established GCs upon loss of MIZ1’s transcriptional activity could be attributed to multiple possibilities, i) enhanced differentiation of IgG1^+^ GC B cells into PCs or MBCs, ii) impaired CSR of IgM to IgG1 early after immunization, and iii) a deficiency in the positive selection of IgG1^+^ GC B cells.

To investigate the first possibility, we employed flow cytometry to assess PC populations and observed a significant decrease in their number in MIZ1^KO^ compared to Ctrl (**Figure 2D**). To further examine the impact on PCs in an isotype-specific manner, we evaluated IgG1 and IgM anti-HEL antibody titers. We found a significant reduction in anti-HEL IgG1 antibody titers in MIZ1^KO^ mice compared to Ctrl. In contrast, the reduction in anti-HEL IgM antibody titers was modest (**Figure 2E**). These findings indicate that the production of IgG1^+^ PCs is diminished following the deletion of MIZ1’s POZ domain. Similarly, the numbers of IgG1^+^ MBCs were significantly reduced in MIZ1^KO^ mice compared Ctrl (**Figure S2B**). Thus, the decreased number of IgG1^+^ GC B cells in MIZ1^KO^ mice was not due to an increased GC output.

We proceeded to investigate whether CSR from IgM to IgG1 was affected in MIZ1^KO^ mice. As CSR occurs prior to the formation of GCs^22^, we examined the number of IgG1^+^ activated B cells during the early stages of the immune response, i.e., before GC formation (**Figure 3A**). By gating on IgD-negative eYFP^+^ non-GC B cells, we identified activated B cells and analyzed the IgG1^+^ and IgM^+^ subsets within this population (**Figure 3B**). We found that the fraction of IgG1^+^ and IgM^+^ activated B cells was comparable between MIZ1^KO^ and Ctrl mice (**Figure 3C**). These results indicate that the decreased number of IgG1^+^ B cells in established GCs in MIZ1^KO^ mice is not due to impaired CSR from IgM to IgG1. To further confirm this, we examined the amplification of IgHγ1-germline transcripts (GLTs) as a marker of CSR from FACS-purified naïve follicular B cells, activated B cells, and GC B cells^77^. Consistent with previous findings^22^, the levels of IgHγ-GLTs were highest in activated B cells, indicating active pre-GC CSR, and these levels were not reduced in MIZ1^KO^ mice compared to the Ctrl (**Figure S3A**). Taken together, these data suggest that CSR is not impaired in MIZ1^KO^ mice.

**Figure. 3.**
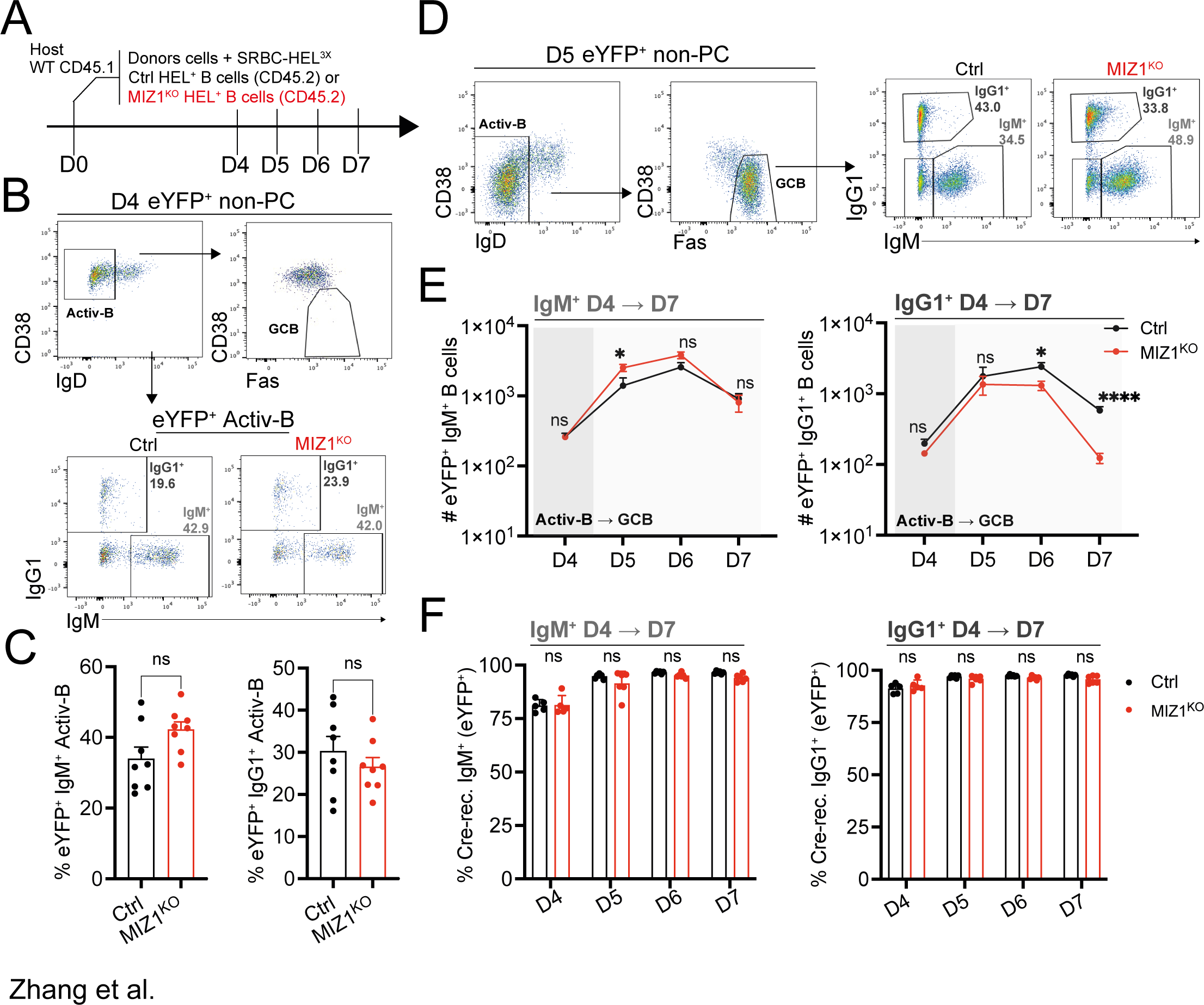
MIZ1 is required for IgG1^+^ B cell accruement in GCs. **(A)** SWHEL experiment schematics. **(B)** Gating strategy for donor-derived activated pre-GC B cells, IgG1^+^ and IgM^+^ B cell subsets. **(C)** Graphs displaying cumulative data of percentages of activated IgG1^+^ and IgM^+^ B cells within total reporter positive pre-GC activated B cells. **(D)** Gating strategy for donor-derived early GC B cells, IgG1^+^ and IgM^+^ GC B cell subsets. **(E)** Dynamic changes of the relative cell numbers (numbers per 10^6^ splenocytes) of reporter positive IgM^+^ and IgG1^+^ activated B cells (D4), and GC B cells (D5-7). **(F)** Dynamic changes of the percentages of eYFP^+^ cells within IgM^+^ (left) and IgG1^+^ (right) activated pre-GC B cells (D4) and GC B cells (D5-7). Each symbol (C: Ctrl n = 8, MIZ1^KO^ n = 8; E: day 4 Ctrl n = 8, MIZ1^KO^ n = 8; day 5 Ctrl n = 6, MIZ1^KO^ n = 6; day 6 Ctrl n = 6, MIZ1^KO^ n = 6; day 7 Ctrl n = 8, MIZ1^KO^ n = 8; F : day 4 – day 7 Ctrl n = 6, MIZ1^KO^ n = 6) represents an individual mouse; small horizontal lines show mean and SEM. Data in (C and E) is from two independent experiment–. Data in (F), D4 - 6 is from two independent experiments, day 7 from three independent experiments. *, P ≤ 0.05; **, P ≤ 0.01; ***, P ≤ 0.001; ****, P ≤ 0.0001 (multiple t test in (A-D). Two-way Anova ns, not significant.)

As the impact of MIZ1’s transcriptional activity on pre-GC IgG1^+^ B cells on day 4 was not evident (**Figure 3C**), it was crucial to determine when the defect in IgG1^+^ GC B cell accruement occurred. To address this, we monitored the dynamic changes in the numbers of IgG1^+^ GC B cells over time. Notably, the numbers of IgG1^+^ GC B cells were comparable between Ctrl and MIZ1^KO^ mice when the GCs were just formed on day 5 (**Figure 3D**, **E**). However, starting from day 6 after immunization, there was a noticeable reduction in the numbers of IgG1^+^ GC B cells in MIZ1^KO^ mice, which became highly pronounced by day 7 (**Figure 3E**). This pattern was also observed for GC B cells that had undergone CSR to other isotypes beyond IgG1 (**Figure S3B**). In contrast, non-switched IgM^+^ GC B cells displayed no defects throughout the observation period (**Figure 3E**).

To rule out the possibility that the specific reduction observed in IgG1^+^ GC B cells was due to Cγ1-cre being expressed only in IgG1^+^ GC B cells, we examined the efficiency of cre-mediated recombination using the eYFP reporter in pre-GC B cells and GC B cells. At all the time-points analyzed, the fraction of eYFP^+^ cells within both IgM^+^ and IgG1^+^ subsets were found to be identical between MIZ1^KO^ and Ctrl mice, indicating that no counterselection or preferential recombination had occurred (**Figure 3F**). Further IgM^+^ and IgG1^+^ GC B cells were nearly 100% eYFP^+^ (**Figure 3F**). This data demonstrates that Cγ1-cre mediated recombination is equally effective in IgM^+^ and IgG1^+^ cells^71^.

### MIZ1 safeguards IgG1^+^ GC B cells during positive selection

To investigate the role of MIZ1 in the positive selection of IgG1^+^ GC B cells, we examined cell proliferation through EdU incorporation. We observed at different time points post-immunization that the proportion of EdU^+^ cells among both IgG1^+^ and IgM^+^ GC B cells was identical between MIZ1^KO^ and Ctrl (**Figure 4A** and **B**). These data suggested that loss of MIZ1’s transcriptional activity did not impact GC B cell proliferation. We then assessed apoptosis by staining for active-Caspase3 (aCasp3^+^), an intracellular marker of cell death. No difference was observed in cell death between Ctrl and MIZ1^KO^ in pre-GC B cells and early GC B cells on day 4 and 5 (**Figure 4C** and **D**). While a modest increase in cell death was observed in IgM^+^ GC B cells in MIZ1^KO^ compared to Ctrl, a pronounced increase was noted in IgG1^+^ GC B cells in MIZ1^KO^ as early as day 6 post-immunization, becoming markedly evident by day 7 (**Figure 4C** and **D**).

**Figure. 4.**
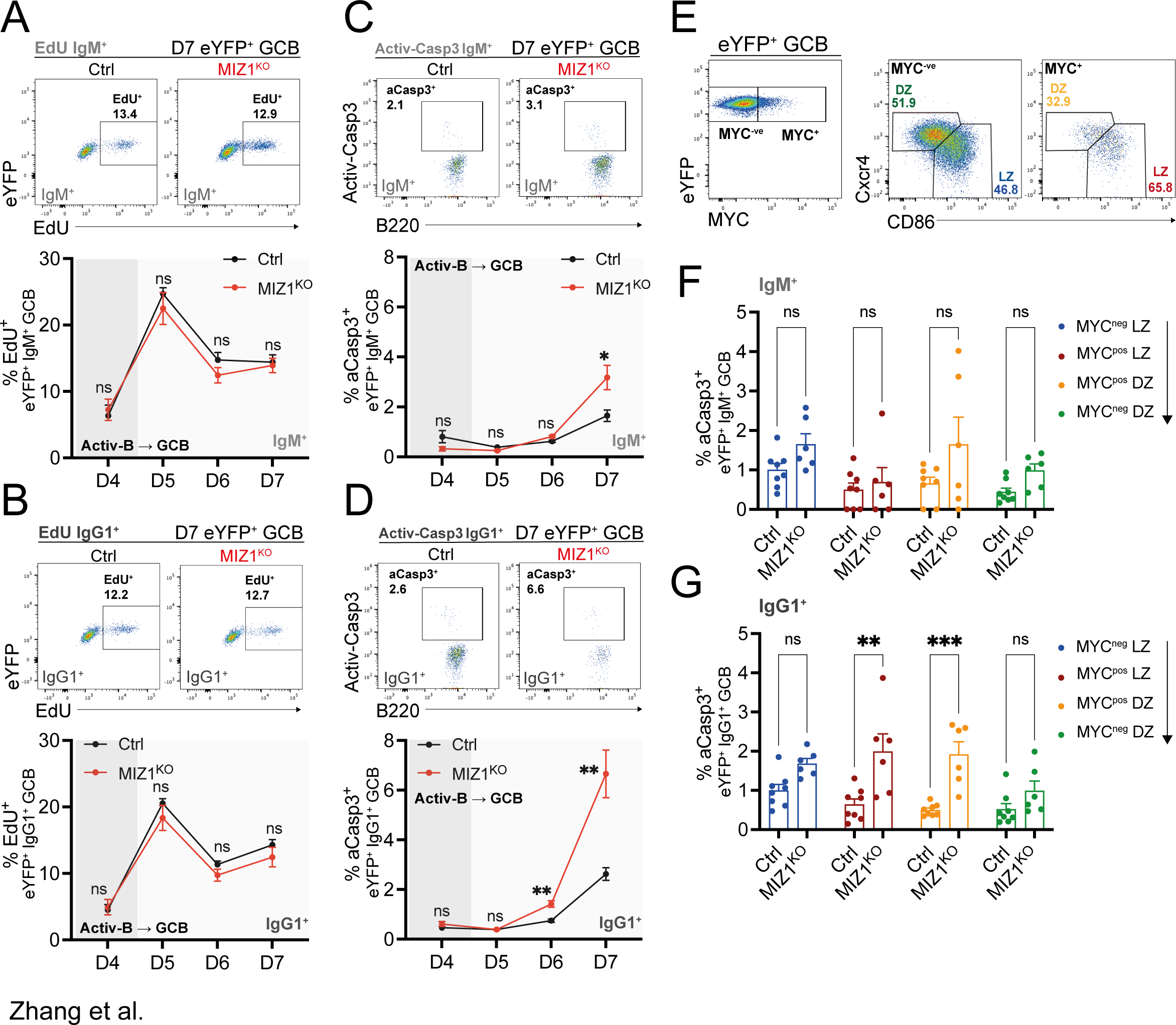
MIZ1 safeguards IgG1^+^ GC B cells during positive selection. SWHEL experiment schematics and gating strategy for IgM^+^ and IgG1^+^ B cells were shown in Fig. 3. **(A)** Top, representative gating strategy for EdU^+^ proliferated IgM^+^ GC B cells (D7). Bottom, dynamic percentages of EdU^+^ cells within IgM^+^ activated pre-GC B cells (D4) and GC B cells (D5-7). **(B)** Top, representative gating strategy for EdU^+^ proliferated IgG1^+^ GC B cells (D7). Bottom, dynamic percentages of EdU^+^ cells within IgG1^+^ activated pre-GC B cells (D4) and GC B cells (D5-7). **(C)** Top, representative gating strategy for active-Caspase3^+^ apoptotic IgM^+^ GC B cells (D7). Bottom, dynamic percentages of aCaspase3^+^ cells within IgM^+^ activated pre-GC B cells (D4) and GC B cells (D5-7). **(D)** Top, representative gating strategy for active-Caspase3^+^ apoptotic IgG1^+^ GC B cells (D7). Bottom, dynamic percentages of aCaspase3^+^ cells within IgG1^+^ activated pre-GC B cells (D4) and GC B cells (D5-7). (E) Gating strategy for MYC^neg^LZ, MYC^pos^LZ, MYC^pos^DZ, MYC^neg^DZ four consecutive GC B cell populations. (F) Normalized percentages of aCaspase3^+^ cells within each of the four consecutive IgM^+^ GC B cell populations. To enable comparison of readings across different experiments, percentages of aCaspase3 were normalized. Firstly, the average percentage of aCasp3^+^ cells in the MYC^neg^LZ cells from the control (Ctrl) group was calculated for each individual experiment. Normalization was then performed by dividing each percentage of aCasp3^+^ cells from both groups by this average number. **(G)** Normalized percentages of aCaspase3^+^ cells within each of the four consecutive IgG1^+^ GC B cell populations. Normalization was done in the same way as described in (F). Each symbol (A-B: D4 – D6 Ctrl n = 6, MIZ1^KO^ n = 6; D7 Ctrl n = 9, MIZ1^KO^ n = 9; C-D: D4-6 Ctrl n = 3, MIZ1^KO^ n = 3; D7 Ctrl n = 8, MIZ1^KO^ n = 8; F-G: Ctrl n = 8, MIZ1^KO^ n = 6) represents an individual mouse; small horizontal lines show mean and SEM. Data in (A, B) is representative from two to three experiments. Data in (C, D), D4-6 is from one experiment, D7 is from two independent experiments. Data in (E, F) is from two independent experiments. *, P ≤ 0.05; **, P ≤ 0.01; ***, P ≤ 0.001; ****, P ≤ 0.0001; Multiple t test in (A-D), two-way ANOVA in (E, F)). ns, not significant.

To gain a deeper understanding of the dynamics of GC B cells, we divided these cells into four consecutive subpopulations^24^: MYC^neg^ LZ → MYC^pos^ LZ → MYC^pos^ DZ → MYC^neg^ DZ (**Figure 4E**). This GC B cell subdivision allowed us to examine the role of MIZ1 in greater detail. Remarkably, the loss of MIZ1’s transcriptional activity had the most significant impact on IgG1^+^ GC B cells in the MYC^pos^ LZ and MYC^pos^ DZ subpopulations. In these subpopulations, we observed a more than three-fold increase in the proportion of apoptotic cells, while no difference was observed for IgM^+^ GC B cells (**Figure 4F** and **G**). This finding highlights the specific role of MIZ1 in safeguarding the survival of IgG1^+^ GC B cells during positive selection.

### MIZ1 regulates the anti-apoptotic factor TMBIM4

To uncover the mechanism underlying the role of MIZ1 in the survival of IgG1^+^ GC B cells during positive selection, we performed RNA sequencing on FACS-purified LZ GC B cells from MIZ1^KO^ and Ctrl. We identified differentially expressed genes (DEGs) using stringent criteria of <0.01 adjusted p-value and >2-fold change. Among the DEGs, we observed reduced expression in MIZ1^KO^ compared to Ctrl of previously reported MIZ1 target genes, including *Rpl22*, *Rscr1*, and *Exoc2*^78–80^ (**Figure 5A**).

**Figure. 5.**
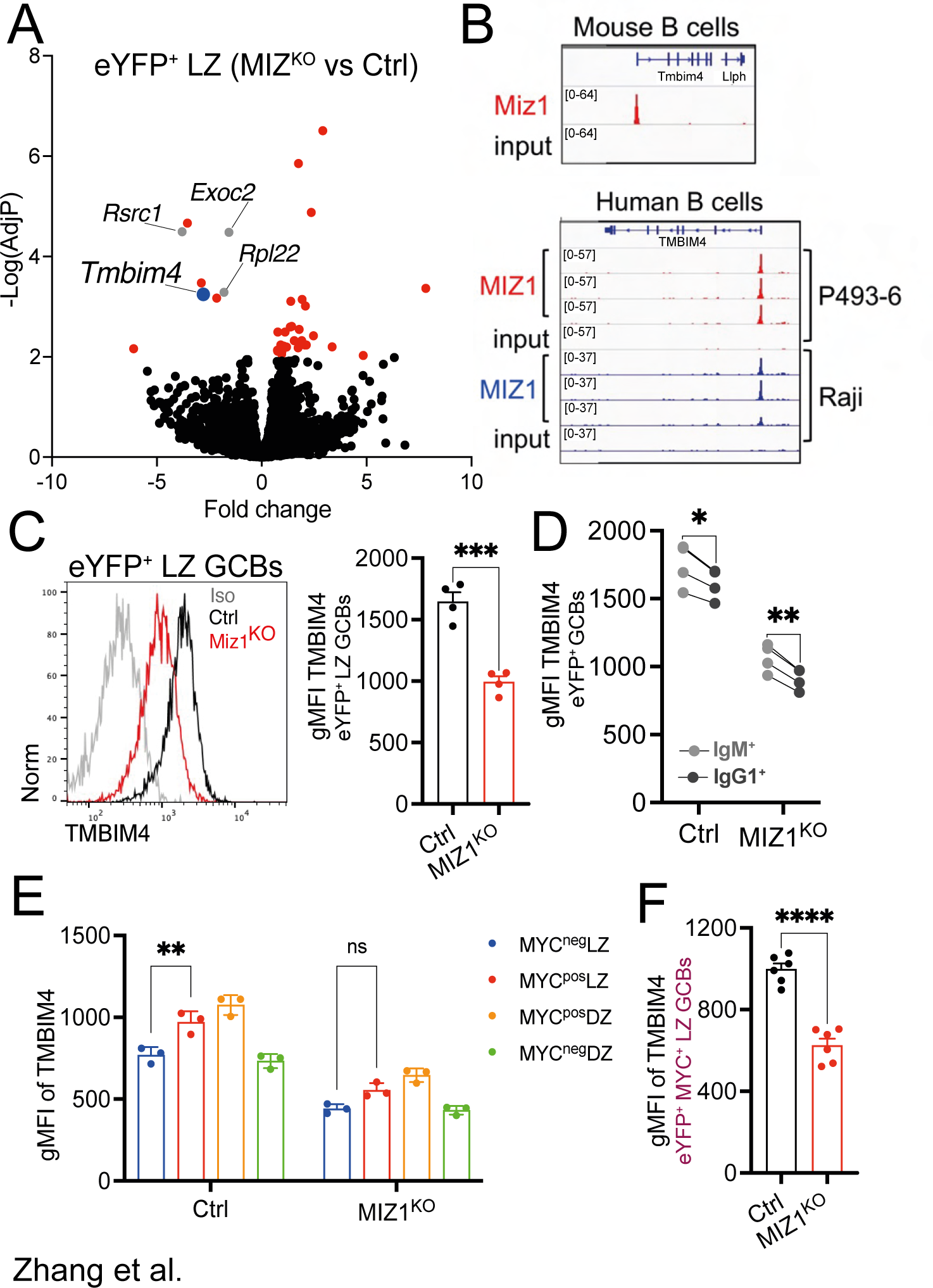
MIZ1 regulates the anti-apoptotic factor TMBIM4. **(A)** Volcano plot for the fold change in gene expression and adjusted P value (-Log(AdjP)) between LZ GC B cells from MIZ1^KO^ mice versus LZ GC B cells from control mice. Colored genes (Red, Blue, and Gray) have a -Log(AdjP) > 2. Genes in Gray are named and are reported MIZ1’s target genes. Blue identifies *Timbim4*. **(B)** Top, MIZ1 ChIP-seq data in mouse primary B cells for binding into the promoter region of the *Tmbim4* gene (data reanalyzed from GSE98419). Bottom, MIZ1 CHIP-seq data in human B cell lines (P493-6, Raji) for binding into the promoter region of the *TMBIM4* gene. **(C-F)** Spleens were collected from CD45.1^+^ recipient mice after 6 days of SRBCs-HEL3x immunization and analyzed by flow cytometry. **(C)** Left, representative plot for TMBIM4 expression levels in LZ GC B cells by intracellular stain. Right, representative data for TMBIM4 expression levels in LZ GC B cells, measured by gMFI. **(D)** Paired analysis for the expression of TMBIM4 in IgM^+^ and IgG1^+^ GC B cells derived from MIZ1^KO^ and control mice, measured by gMFI. **(E)** Representative data for TMBIM4 expression levels in MYC^neg^ LZ, MYC^pos^ LZ, MYC^pos^ DZ, MYC^neg^ DZ consecutive GC B cell populations from control and MIZ1^KO^ mice. **(F)** Comparison of TMBIM4 expression levels in Myc^pos^LZ GC B cells derived from MIZ1^KO^ and control mice. Each symbol (C-D: Ctrl n = 4, MIZ1^KO^ n = 4; E: Ctrl n = 3 MIZ1^KO^ n = 3; F: Ctrl n = 6, MIZ1^KO^ n = 6) represents an individual mouse; small horizontal lines show mean and SEM. Data in (C-D) is one representative of four independent experiments. Data in (D) is one representative of two independent experiments. **, P ≤ 0.01; ***, P ≤ 0.001; ****, P ≤ 0.0001 (unpaired two-tailed Student’s t test in (C, F); multiple paired t test in (E); Two-way ANOVA (D, F)).

We also found that the expression of the gene encoding the anti-apoptotic protein TMBIM4^41^ was significantly downregulated in MIZ1^KO^ compared to Ctrl. To investigate if the decreased expression of *Tmbim4* in MIZ1^KO^ mice was directly linked to the loss of MIZ1’s transcriptional activity, we conducted chromatin immunoprecipitation sequencing (ChIP-seq) of MIZ1 in mouse and human B cells. These data revealed the specific enrichment of MIZ1 at the promoter region of *Tmbim4* compared to input control in mouse B cells. Moreover, this finding was corroborated by the binding of MIZ1 to the *TMBIM4* promoter region in two human B cell lines (**Figure 5B**). These results suggest a direct role for MIZ1 in controlling TMBIM4 expression.

To evaluate the impact of impaired *Tmbim4* transcription on protein expression, we performed TMBIM4 staining on GC B cells. We observed a significant decrease in TMBIM4 levels in the LZ GC B cells of MIZ1^KO^ compared to Ctrl (**Figure 5C**). The loss of MIZ1’s transcriptional activity led to reduced TMBIM4 expression in both IgG1^+^ and IgM^+^ GC B cells. However, the expression of TMBIM4 was less impacted in IgM^+^ GC B cells from MIZ1^KO^ mice compared to their IgG1^+^ counterparts (**Figure 5D**). The analysis of the expression pattern of TMBIM4 in GC B cells revealed a reduction across all four subpopulations (MYC^neg^ LZ → MYC^pos^ LZ → MYC^pos^ DZ → MYC^neg^ DZ) in the absence of MIZ1’s transcriptional activity (**Figure 5E**). However, in the Ctrl, there was a significant induction of TMBIM4 from MYC^neg^ LZ → MYC^pos^ LZ i.e., at the positive selection stage. Notably, this induction of TMBIM4 was lost in MIZ1^KO^ (**Figure 5E**). More specifically, there was roughly a 40% reduction in TMBIM4 levels in MYC^pos^ LZ cells of MIZ1^KO^ compared to Ctrl (**Figure 5F**). These data suggest that MIZ1 directly regulates *Tmbim4/TMBIM4* expression, a newly identified target gene that encodes an anti-apoptotic protein.

### MIZ1 controls Ca^2+^ mobilization during positive selection

TMBIM4 is a member of an ancient protein family, known for its anti-apoptotic functions^40^. This family of proteins is highly conserved across different species and are primarily localized on the Golgi apparatus and endoplasmic reticulum (ER) membranes^41,81,82^. Previous studies *in vitro* using epithelial cancer cells showed that TMBIM4 regulates the release of ER Ca^2+^ via its interaction with IP3R^43^ (**Figure 6A**). Given the control of TMBIM4 expression by MIZ1 we investigated whether the regulation of Ca^2+^ release was altered in MIZ1^KO^. First we generated iGB cells using both Ctrl and MIZ1^KO^ naïve B cells and performed analysis when >80% of cells were immunoglobulin isotype class switched^48^. Upon stimulation of iGB cells with anti-Ig F(ab’)2, we observed exacerbated Ca^2+^ mobilization in MIZ1^KO^ cells compared to Ctrl, with sustained high Ca^2+^ levels in MIZ1^KO^ cells throughout the observation period (**Figure 6B** and **Supplemental Video 2**). To further investigate the role of MIZ1 in Ca^2+^ regulation, we performed *ex vivo* stimulation of freshly isolated GC B cells (**Figure S4A**). There was a slight trend towards higher Ca^2+^ levels in MIZ1^KO^ IgM^+^ GC B cells compared to Ctrl; however, the difference did not reach statistical significance (**Figure 6C**). In contrast, we found a highly exacerbated and significant increase in cytosolic Ca^2+^ levels in IgG1^+^ GC B cells of MIZ1^KO^ compared to Ctrl (**Figure 6C**). Notably, the loss of MIZ1’s transcriptional activity did not change surface BCR levels in GC B cells (**Figure S4B**). To further characterize Ca^2+^ release in an isotype-specific manner, we compared IgM^+^ and IgG1^+^ GC B cells from each individual mouse. This analysis revealed that physiologically, IgG1^+^ GC B cells exhibited a naturally increased Ca^2+^ mobilization compared to IgM^+^ GC B cells, and that this difference became exacerbated in MIZ1^KO^ (**Figure 6D**). This is consistent with data showing enrichment for gene signatures associated with Ca^2+^ entry, release, and import in IgG1^+^ versus IgM^+^ GC B cells^20^ (**Figure S4C**). We conclude that the transcriptional mechanism composed by the MIZ1-TMBIM4 axis primarily controls Ca^2+^ mobilization IgG1^+^ GC B cells.

**Figure. 6.**
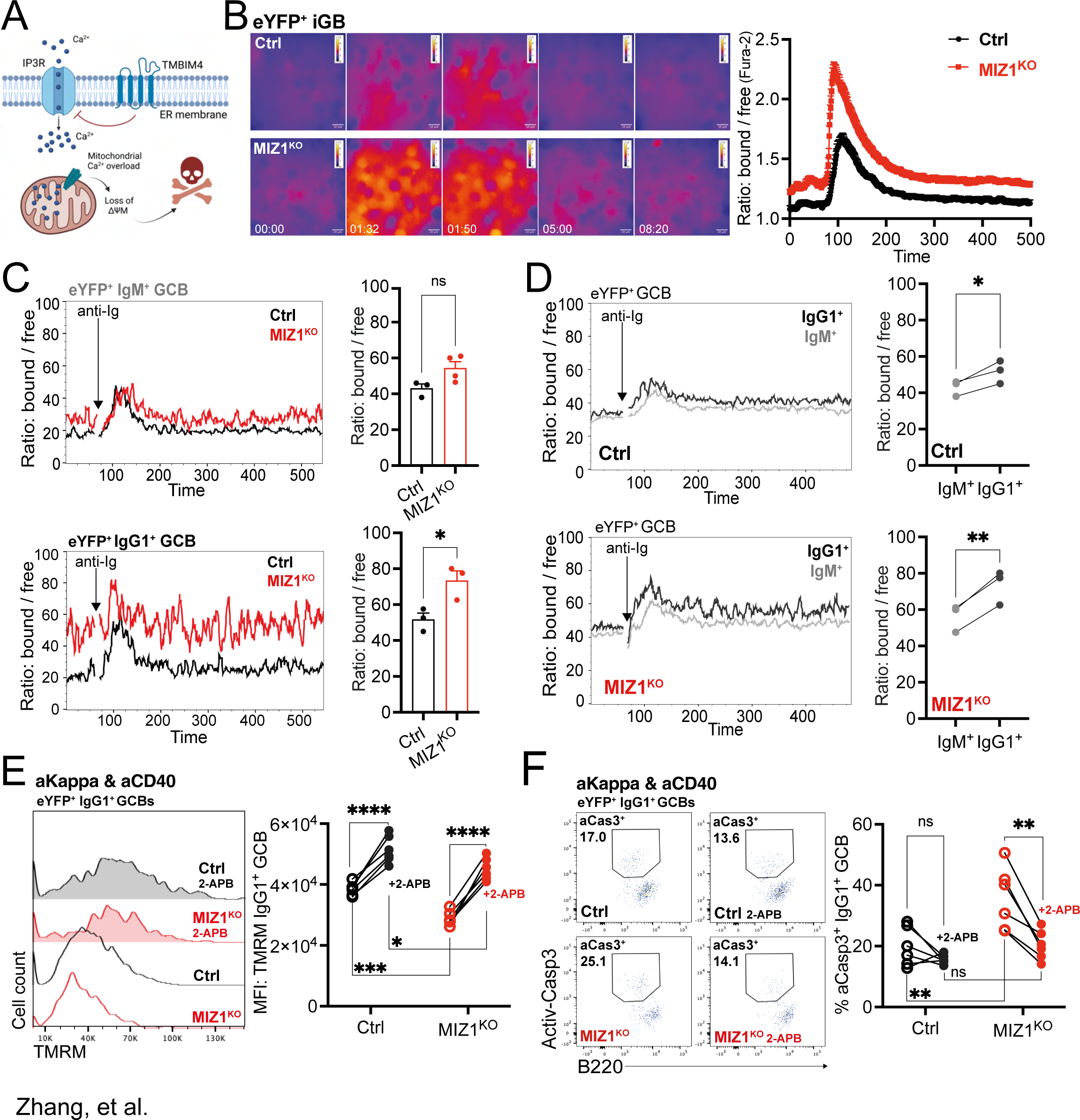
MIZ1 controls Ca^2+^ mobilization during positive selection. **(A)** Schematic diagram exemplifying the ER membrane, and TMBIM4 inhibition of Ca^2+^ release through modulation of IP3R. Created with BioRender.com. **(B)** Measurement of Ca^2+^ flux by live cell imaging. Fura-2 stained iGB cells were stimulated with anti-Ig antibody. Ratiometric images were generated by ImageJ. Left: time-point dynamics of Ca^2+^ flux before (00:00) and after stimulation (01:32, 01:50, 05:00 and 08:20). Right, analysis of ratio: bond / free reflecting cytoplasmic Ca^2+^ levels. Pooled data from 25 cells for each genotype. **(C-F)** Splenocytes were collected from control and MIZ1^KO^ mice after 8-9 days of 1x 10^9^ SRBCs immunization. **(C)** Left, representative Ca^2+^ flux dynamics calcium flux for Indo-1-stained IgG1^+^ and IgM^+^ GC B cells; right, graphs of cumulative data for peak cytoplasmatic levels of Ca^2+^ after stimulation. control is black line, MIZ1^KO^ is red line. Cells were stained with Indo-1 and *ex-vivo* stimulated with anti-Ig antibody (10 ug/ml) during sample acquiring for calcium flux analysis. **(D)** Left, representative Ca^2+^ flux between IgG1^+^ (dark grey) and IgM^+^ (light grey) GC B cells in each individual mouse of the same genotype. Right, peak cytoplasmatic levels of Ca^2+^ after stimulation. Top, control; bottom, MIZ1^KO^. **(E-F)** RBCs depleted splenocytes were *in vitro* stimulated with F(ab’)2 anti-Kappa (1 ug/ml) and (or) anti-CD40 (0.5 ug/ml) antibody in the presence or without 2APB (100 mM) for 3 hours and stained for TMRM or active-Caspase3 for flow cytometry analysis. **(E)** Impact of IP3R inhibitor 2-APB on ΔΨ_M_. Left, representative TMRM levels from *ex vivo* stimulated IgG1^+^ GC B cells treated (filled plots) or not (single line) with 2-APB. Right, graph displaying cumulative data paired analysis. MFI of TMRM is shown. **(F)** Impact of IP3R inhibitor 2-APB on apoptosis. Left, representative gating strategy for active-Caspase3^+^ (aCasp3^+^) in *ex vivo* stimulated IgG1^+^ GC B cells treated (filled plots) or not (single line) with 2-APB. Right, graph displaying cumulative data of percentages of active-Casepase3^+^ IgG1^+^ GC B cells. Each symbol (C-D: Ctrl n = 3 or 4, MIZ1^K^^O^ n = 3; E-F: Ctrl n = 6, MIZ1^KO^ n = 6) represents an individual mouse; small horizontal lines show mean and SEM. Data in (B) is representative of four experiments. Data in (C-D) is representative from 3 independent experiments. Data in (E-F) is representative of two experiments. *, P ≤ 0.05; **, P ≤ 0.01 (unpaired two-tailed Student’s t test in (C,); paired t test in (D); Two-way ANOVA (E, F). ns, not significant.

### IP3R inhibition rescues ΔΨM and protects MIZ1^KO^ IgG1^+^ GC B Cells

To gain further insights into the impact of Ca^2+^ mobilization in IgG1^+^ GC B cells during positive selection, we conducted *ex vivo* experiments using freshly isolated GC B cells stimulated with anti-Igκ and anti-CD40, known to induce robust MYC expression and to mimic positive selection^17^ (**Figure S4D**). We assessed the extent of apoptosis by quantifying the fraction of cells positive for active Caspase-3 (aCasp3^+^). Remarkably, the stimulation with anti-Igκ plus anti-CD40 significantly reduced cell death of IgG1^+^ GC B cells from Ctrl, compared to anti-Igκ alone. However, and in contrast, such a rescue was significantly impaired in MIZ1^KO^, with the apoptotic rate being nearly twice as high as in Ctrl (**Figure S4E**). These data are consistent with the *in vivo* observations (**Figure 4D** and **G**) and emphasizes the increased the vulnerability of MIZ1^KO^ IgG1^+^ GC B cells to cell death, despite TFH-help.

Apoptosis can be induced by an increase in intracellular calcium levels and is characterized by mitochondrial outer membrane permeabilization (MOMP), leading to the release of pro-apoptotic factors^83^. MOMP is directly linked to the loss of mitochondrial membrane potential (ΔΨM)^84^ (**Figure 6A**). To assess the mitochondrial function in GC B cells, we measured the maximum ΔΨM in both Ctrl and MIZ1^KO^ cells. Anti-Igκ plus anti-CD40 *ex vivo* stimulated MIZ1^KO^ IgG1^+^ GC B cells exhibited a significant reduction in their maximum ΔΨM compared to Ctrl, indicating impaired mitochondrial function (**Figure 6E**). To directly examine if the loss of ΔΨM and increased cell death observed in MIZ1^KO^ IgG1^+^ GC B cells was a consequence of uncontrolled IP3R-mediated Ca^2+^ release, we used the IP3R antagonist 2-APB^85,86^. Inhibition of IP3R Ca^2+^ release by 2-APB effectively restored the maximum ΔΨM of MIZ1^KO^ IgG1^+^ GC B cells (**Figure 6E**) and curtailed cell death (**Figure 6F**). The impact of MIZ1^KO^ on ΔΨM and cell death of IgM^+^ GC B cells was relatively mild (**Figure S4F, G**). These data indicated that exacerbated Ca^2+^ release from IP3R in MIZ1^KO^ IgG1^+^ GC B cells underlie cell death.

### Restoring TMBIM4 rescues MIZ1^KO^ IgG1^+^ GC B cell positive selection

To determine whether exacerbated Ca^2+^ release in MIZ1^KO^ IgG1^+^ GC B cells was directly linked to impaired TMBIM4 expression, we conducted experiments with iGB cells to restore TMBIM4 levels and to analyze Ca^2+^ mobilization. We generated a retroviral vector that overexpressed TMBIM4 by inserting a *Tmbim4-T2A-RFP* expression cassette downstream of the 5’ long terminal repeat (LTR). As a control, we constructed a retroviral vector containing only *RFP* expression. By detecting RFP-positive retrovirally transduced iGB cells, we found that restoration of TMBIM4 effectively reduced Ca^2+^ mobilization in MIZ1^KO^ (**Figure 7A**). Thus, TMBIM4 restoration compensated the loss of MIZ1’s transcriptional activity.

**Figure. 7.**
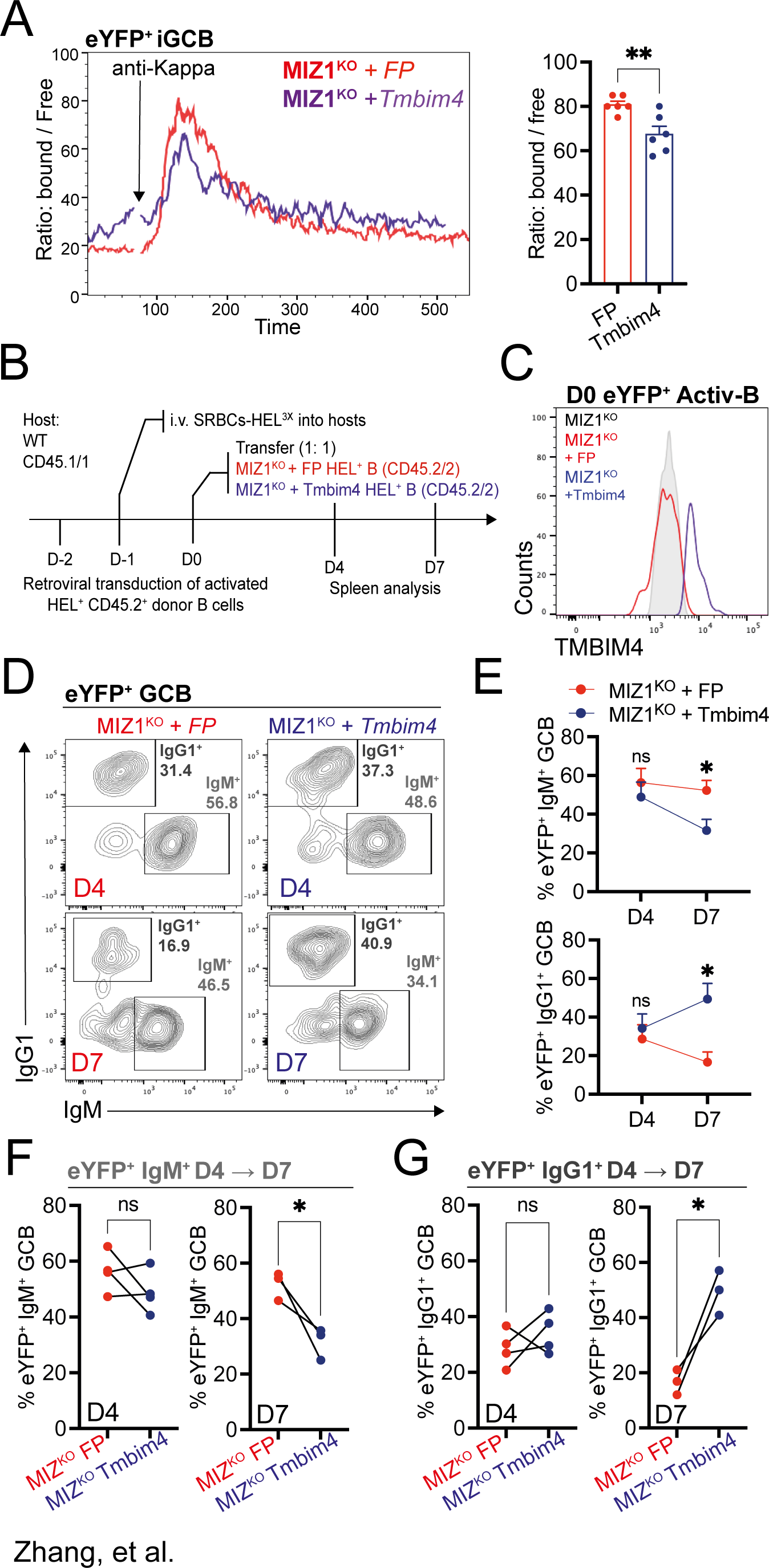
Restoring TMBIM4 rescues MIZ1^KO^ IgG1^+^ GC B cell positive selection. **(A)** Left, representative Ca^2+^ flux dynamics for Indo-1 stained iGB cells upon restoration (MIZ1^KO^ + *Tmbim4*) or not (MIZ1^KO^ + *FP*) of TMBIM4 in MIZ1^KO^ cells. Right, cumulative data for Ca^2+^ flux showing peak cytoplasmatic levels after stimulation. **(B)** Schematics of the experimental workflow for the restoration of TMBIM4 expression *in vivo*. **(C)** Representative plots of *ex vivo* GC B cells displaying restoration (MIZ1^KO^ + *Tmbim4*) or not (MIZ1^KO^ + *FP*) of TMBIM4. **(D)** Gating strategy for IgG1^+^ and IgM^+^ GC B cell subsets generated from donor-derived reporter positive B cells. **(E)** Dynamic changes of the percentages of IgG1^+^ and IgM^+^ B cell subsets in retrovirally transduced GC B cell on day 4 and day 7. **(F)** Representative data for IgM^+^ B cell percentages within GC B cells upon restoration (MIZ1^KO^ + *Tmbim4*) or not (MIZ1^KO^ + *FP*) of Tmbim4 in MIZ1^KO^ cells. Left, day 4 after cell transfer; right, day 7 after cell transfer. **(G)** Representative data for IgG1^+^ B cell percentages within GC B cells upon restoration (MIZ1^KO^ + *Tmbim4*) or not (MIZ1^KO^ + *FP*) of Tmbim4 in MIZ1^KO^ cells. Left, day 4 after cell transfer; right, day 7 after cell transfer. Each symbol (A: MIZ1^KO^ + *FP* n = 6, MIZ1^KO^ + *Tmbim4* n = 6; D-G: day 4 n = 4; day 7 n = 3) represents an individual mouse. Data in (A) is from two independent experiments, data in (D-G) is one representative from two independent experiments. *, P ≤ 0.05; **, P ≤ 0.01 (unpaired two-tailed Student’s t test in (A); multiple t test (E); paired t test (F-G)). ns, not significant.

To examine whether restoring TMBIM4 expression rescues *in vivo* IgG1^+^ GC B cell positive selection in MIZ1^KO^ mice, we performed adoptive transfer experiments. Equal numbers of MIZ1^KO^ HEL^+^ B cells transduced with either the control or the *Tmbim4* expressing vector were transferred into the same recipient mouse according to established protocols^87^. Subsequently, recipient mice were immunized with HEL^3X^ (**Figure 7B**). Flow cytometry analysis allowed us to distinguish between control reporter vector transduced B cells (MIZ1^KO^ + FP) and *Tmbim4* expressing transduced B cells (MIZ1^KO^ + Tmbim4). As expected, MIZ1^KO^ GC B cells carrying the *Tmbim4* expressing vector exhibited increased TMBIM4 expression compared to those carrying the control vector (**Figure 7C**). We examined the transduced MIZ1^KO^ GC B cells at day 4, representing early formed GCs in this system, and at day 7, when GCs had matured (**Figure 7D**). The fraction of MIZ1^KO^ + FP IgG1^+^ GC B cells decreased over time between day 4 and day 7, indicating impaired positive selection of IgG1^+^ GC B cells. In contrast, restoration of TMBIM4 in MIZ1^KO^ (MIZ1^KO^ + Tmbim4) rescued positive selection of IgG1^+^ GC B cells (**Figure 7E**). This was evidenced by a decreased fraction of IgM^+^ and an increased fraction of IgG1^+^ GC B cells at day 7 (**Figure 7F**, **G**). These data demonstrates that the protective mechanism orchestrated by the MIZ1-TMBIM4 axis specifically safeguards IgG1^+^ GC B cell positive selection, thus uncovering a distinct dependency specific to this immunoglobulin isotype on a previously unknown mechanism during GC positive selection.

## Discussion

During an adaptive immune response triggered by infection or vaccination, B cells undergo a remarkable process of refinement. This process involves the transition from IgM to IgG accompanied by an increase in affinity of the BCR towards the antigen, ultimately leading to the production of neutralizing antibodies^1–5^. The focal point of this transformative process is the GC, a specialized microenvironment where affinity maturation and preferential accrual of IgG^+^ over IgM^+^ B cells occurs^13,14,21^. However, it is not known whether the GC positive selection of immunoglobulin isotypes depends variably on transcriptional mechanisms.

In this study, we demonstrate the essential and specific role of MIZ1 in protecting IgG1^+^ GC B cells during positive selection. We uncovered that MIZ1 plays a key role in inducing the expression of TMBIM4, a newly described target, member of an ancient protein family with anti-apoptotic properties^40,41^. TMBIM4 effectively controlled Ca^2+^ release via IP3R^42–45^ downstream of IgG1 BCR. This ensured the survival of IgG1^+^ GC B cells during positive selection, while IgM^+^ GC B cells were largely independent. Conversely, in the absence of the MIZ1-TMBIM4 axis, Ca^2+^ mobilization became exacerbated specifically in IgG1^+^ GC B cells leading to mitochondrial dysfunction and subsequent cell death. Restoration of TMBIM4 expression *in vivo* successfully rescued the positive selection of MIZ1-deficient IgG1^+^ GC B cells (**Figure S5**). The role of the MIZ1-TMBIM4 axis in IgM^+^ GC B cells appears to be minimal, suggesting a distinctive dependency on this transcriptional mechanism compared to IgG1^+^ GC B cells. Interestingly, the lack of MIZ1 had a lesser impact on TMBIM4 expression in IgM^+^ GC B cells than in their IgG1^+^ counterparts. This suggests the presence of alternative pathways that could be modulating TMBIM4 expression in IgM^+^ GC B cells and/or other mechanism regulating Ca^2+^ mobilization following the activation of the IgM BCR.

This study marks the first identification of MIZ’s critical role in controlling Ca^2+^ release, and it affirms the *in vivo* importance of the anti-apoptotic factor, TMBIM4. Importantly, this research unveils a dependency, specific to an immunoglobulin isotype, on a hitherto unidentified transcriptional mechanism during positive selection in GCs. Although the study primarily focused on elucidating the role of the MIZ1-TMBIM4 axis in IgG1^+^ GC B cells, which comprise the larger proportion of the IgG pool, we also observed that other class-switched GC B cells (i.e., IgM^neg^IgG1^neg^) were similarly impacted when the axis was disrupted. Over time during GC responses, the lack of MIZ1-TMBIM4 led to a significant decrease in cell numbers of class-switched GC B cells, indicating a broader role for the axis in regulating Ca^2+^-driven mitochondrial dysfunction-induced cell death.

These findings highlight the crucial importance of precise regulation of Ca^2+^ mobilization for the positive selection of IgG GC B cells. Contrary to IgM, BCRs belonging to all IgG subclasses feature an extended cytoplasmic tail^88^. This feature bestows class-switched B cells with enhanced response capabilities^29,33,89^, including increased Ca^2+^ mobilization^30–32^, compared to their unswitched counterparts. Notably, despite seeming like a benefit, the presence of the IgG1 cytoplasmic tail is not required for IgG1^+^ GC B cell positive selection^21^. Considering this intriguing observation alongside insights from the current study, which underscore tight regulation of Ca^2+^ mobilization, it is tempting to speculate that the IgG cytoplasmic tail, might if engaged, adversely impact IgG1^+^ GC B cell the positive selection.

Despite significant advancements in the understanding of BCR-Ca^2+^ signaling, its activity in GC B cells has remained elusive. In naïve B cells, Ca^2+^ signaling serves as a crucial second messenger downstream of BCR, governing multiple intracellular processes and exerting dual effects on cell survival and apoptosis^90^. During the early phase of signaling, antigen engagement with the BCR promotes cell survival^28,91^. However, stimulated B cells may experience a gradual decline in mitochondrial function due to Ca^2+^ accumulation, ultimately leading to cell death unless rescued in time, for example by T cell help^92–94^. In contrast, the current understanding of GC B cell BCR signaling suggests an attenuated and rewired nature^17,34,35^, resulting in compromised Ca^2+^ mobilization compared to naïve B cells^95,96^. Consequently, it remained unclear whether Ca^2+^ can exert dual effects on GC B cell survival and apoptosis, as observed in naïve B cells. Evidence from a transcriptional reporter of BCR signaling (i.e., Nur77 reporter)^36,37^, advanced imaging techniques using high-content^38^ and intravital microscopy^39^ indicate that GC B cell BCR signaling is sufficient to mobilize Ca^2+^. This is potentially related to the observed early survival effects in antigen stimulated GC B cells *in vivo*^28,97^ and with other work in which GC B cells were susceptible to hyperactive BCR signaling induced death^96,98^. To our knowledge these studies have not provided immunoglobulin isotype resolution. However, given the new understanding from the present study on the capacity of IgG1 BCR-mediated Ca^2+^ mobilization to induce mitochondrial dysfunction-mediated cell death, future work is warranted.

The present study suggests that BCR-induced Ca^2+^ mobilization is a life-or-death checkpoint for IgG1^+^ GC B cell positive selection. This could have profound implications for humans, considering the high conservation of TMBIM4 and MIZ1 and of the regulatory role of MIZ1 on TMBIM4 expression in human B cells. The high susceptibility of IgG1^+^ GC B cells to Ca^2+^-mediated cell death following BCR signaling may be advantageous in certain contexts. Within GCs, B cells undergo SHM in response to foreign antigens. However, this process also generates GC B cells that have the potential to react with self-antigens, which poses a risk of compromising self-tolerance and giving rise to autoantibody production^99–103^. To counter this challenge, a rapid and efficient mechanism is necessary to weed out self-reactive GC B cells. The high susceptibility of IgG1^+^ GC B cells to Ca^2+^-mediated cell death can also have detrimental consequences by potentially causing the demise of GC B cells carrying higher-affinity BCRs specific to antigens derived from an infectious agent or vaccination. This underlines the potential significance of actively fine-tuning the MIZ1-TMBIM4 axis during the GC reaction. Future research should explore whether such modulation indeed takes place and assess its implications in autoimmune diseases, infections, and vaccination responses.

## Limitations

During positive selection the MIZ1-TMBIM4 axis limits exacerbated Ca^2+^ mobilization and cell death of IgG1^+^ GC B cells. Although we used multiple immunization conditions, monoclonal and polyclonal systems, it will be relevant to study if and how infectious agents interfere with the activity of the MIZ1-TMBIM4 axis. We have also not explored in detail the role of the MIZ1-TMBIM4 axis beyond IgG1 and IgM GC B cells. Thus, whether this axis is required to prevent cell death of GC B cells carrying other isotypes, namely IgA and IgE remains to be investigated.

## Supporting information

Supplemental Video 2 (CTRL)

Supplemental Video 2 (MIZ1KO)

## Acknowledgments

We thank the members of the Immunity and Cancer laboratory (Francis Crick Institute [FCI], London, UK); O. Bannard (University of Oxford, Oxford, UK), R. Brink (Garvan Institute of Medical Research, Darlinghurst, Australia), V. Tybulewicz (FCI, London, UK) for critical discussions and comments. We thank O. Bannard (University of Oxford, Oxford, UK) for a HEL^3X^-expressing cell line, R. Brink (Garvan Institute of Medical Research, Darlinghurst, Australia) for SWHEL mice, and the FCI scientific platforms (Biological Resource Facility, Flow Cytometry, Histopathology, Light Microscopy, Structural Biology and Advanced Sequencing) for expert advice and technical support. This work was supported by the FCI, which receives core funding from Cancer Research UK (grants CC2078 and CC2085), the UK Medical Research Council (grants CC2078, and CC2085), and the Wellcome Trust (grants CC2078 and CC2085) to D.P. Calado and A. Wack, and Medical Research Council career development award MR/J008060/1 and CRUK [C355/A26819], FC AECC [C355/A26819], AIRC [C355/A26819] to D.P. Calado.

## Supplemental Figure Legends

**Video S1**

Representative three-dimensional imaging of MIZ1 and MYC colocalization in positively selected (MYC^+^) GC B cells. Nucleus was identified with DAPI (blue) staining, and counterstained with MIZ1 (green) and MYC (red). Imaging was generated by Imaris software.

**Figure. S1.**
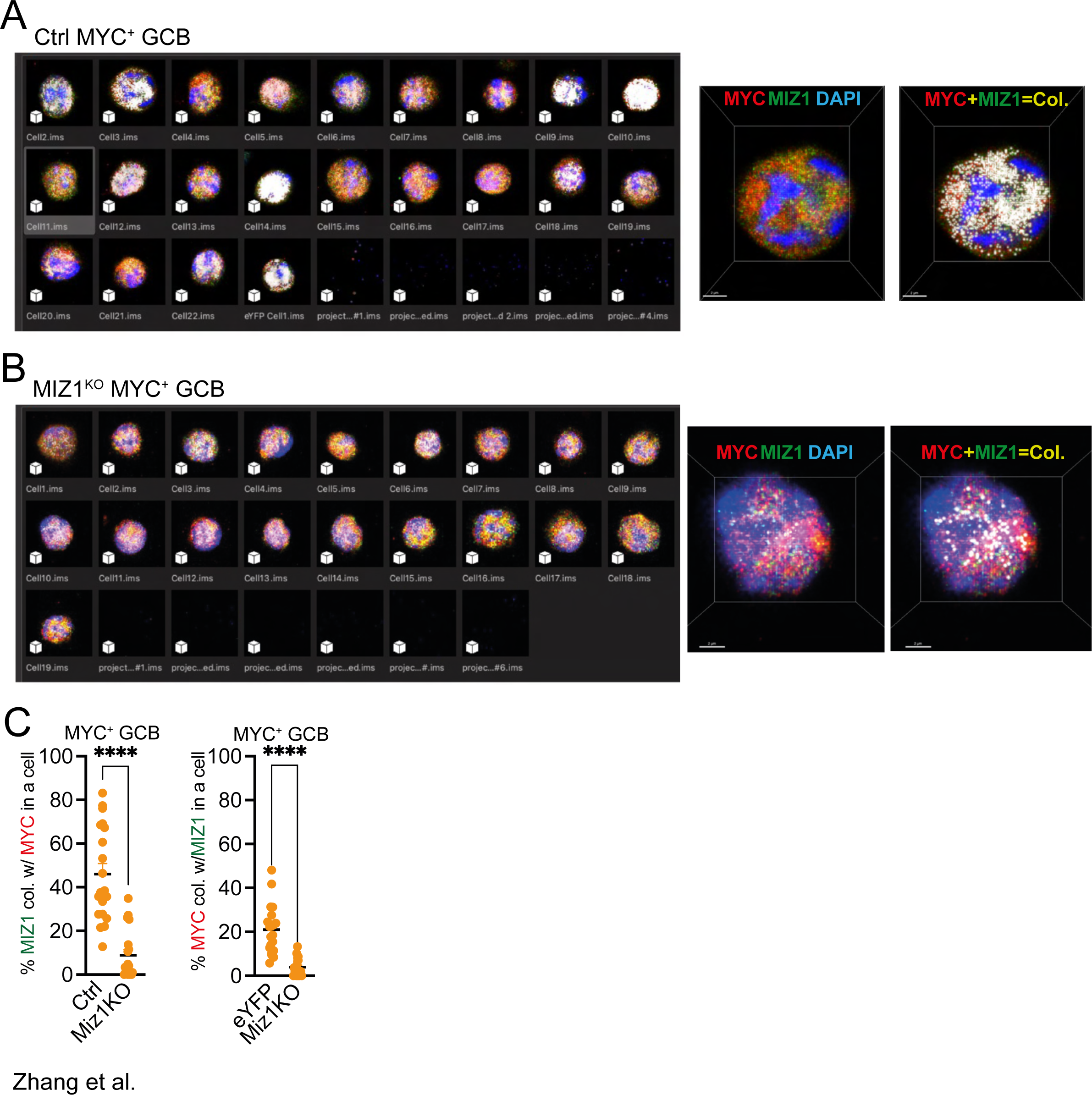
3D image analysis of positively selected GC B cells for MIZ1 and MYC colocalization. **(A)** GC B cells were purified from control mice, and counterstained with DAPI (blue), anti-BCL-6 antibody (grey), anti-MIZ1 antibody (green) and anti-MYC antibody (red). Data analyzed using Imaris software. **(B)** GC B cells were purified from MIZ1^KO^ mice, and counterstained with DAPI (blue), anti-BCL-6 antibody (grey), anti-MIZ1 antibody (green) and anti-MYC antibody (red). Data analyzed using Imaris software. **(C)** Left, percentages of MIZ1 colocalized with MYC; right, percentages of MYC colocalized with MIZ1. Each dot represents one cell, small horizontal line shows mean and SEM. Twenty individual MYC^+^ GC B cells from immunized WT and MIZ1^KO^ mice were displayed and calculated for MYC-MIZ1 colocalization. ****, P ≤ 0.0001 (unpaired two-tailed Student’s t test).

**Figure. S2.**
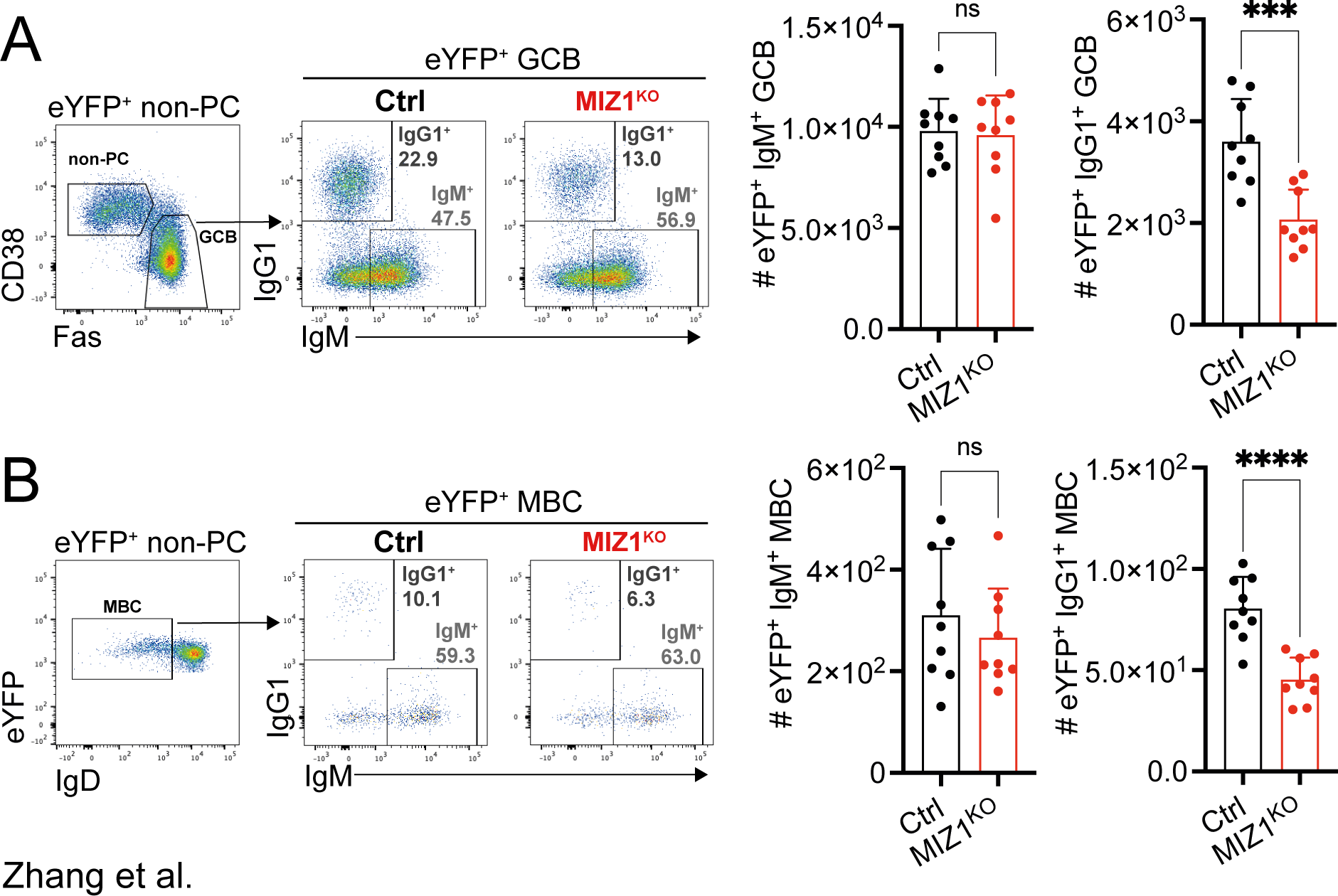
Impaired IgG1^+^ GC B cells and MBCs in MIZ1^KO^ mice. (**A-B**) Control and MIZ1^KO^ mice were i.v. immunized with 1 x 10^9^ SRBCs, and their spleens were collected for flow cytometry analysis after 9 days of immunization. (A) Left, gating strategy for eYFP^+^ GC B cells, IgM^+^ and IgG1^+^ GC B cell subsets. Right, Relative cell numbers (numbers per 10^6^ splenocytes) of IgM^+^ and IgG1^+^ GC B cells in control and MIZ1^KO^ mice. (B) Left, gating strategy for eYFP^+^ MBCs, IgM^+^ and IgG1^+^ memory B cell subsets. Right, Relative cell numbers (numbers per 10^6^ splenocytes) of IgM^+^ and IgG1^+^ MBCs in control and MIZ1^KO^ mice. Each symbol (A-B: Ctrl n = 9, MIZ1^KO^ n = 9) represents an individual mouse; small horizontal lines show mean, and SEM. Data is from three independent experiments. *, P ≤ 0.05; **, P ≤ 0.01; ***, P ≤ 0.001; ****, P ≤ 0.0001 (unpaired two-tailed Student’s t test. ns, not significant.)

**Figure. S3.**
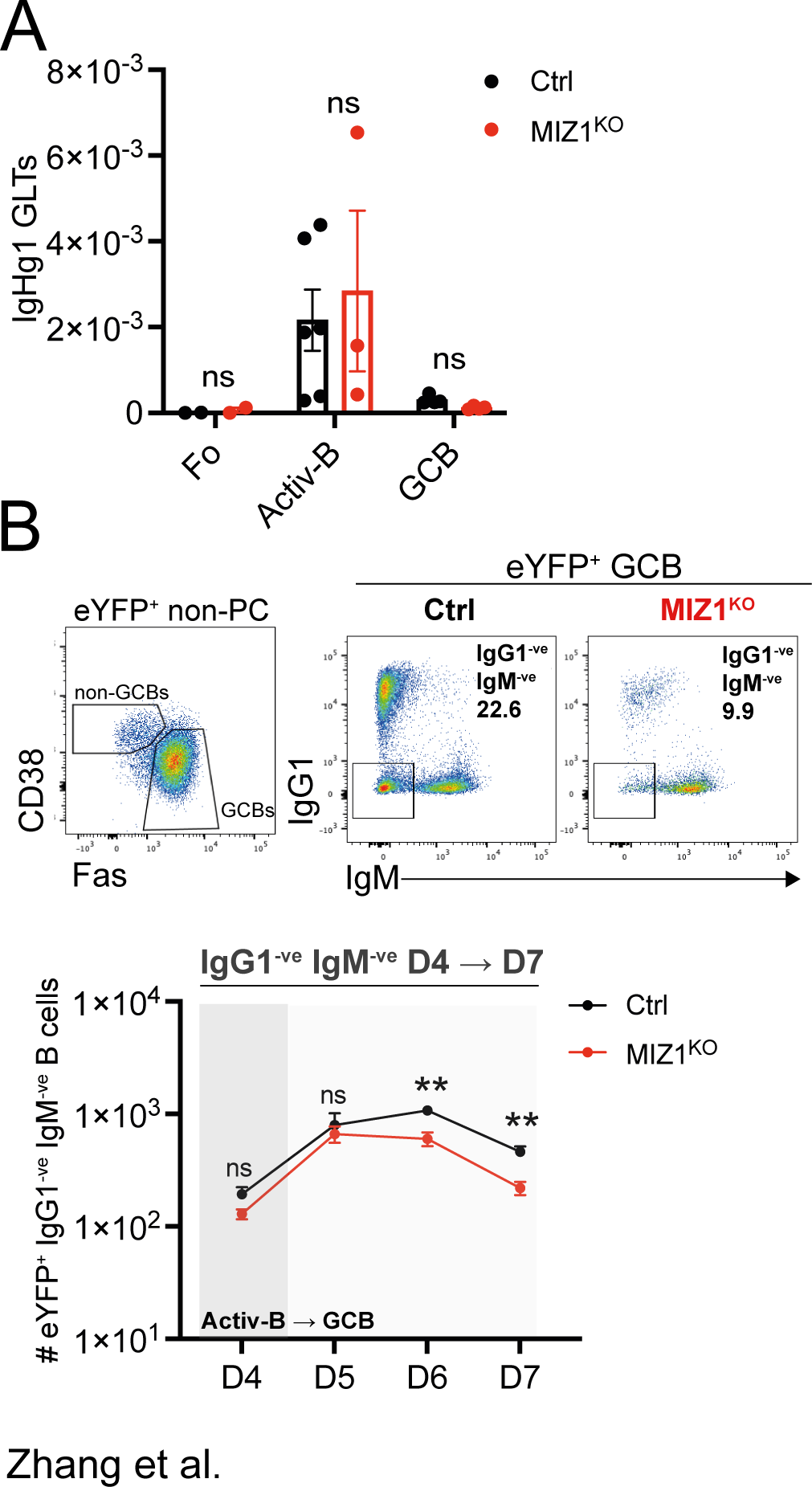
Normal CSR but impaired IgG1^neg^ IgM^neg^ GC B cell accruement *in vivo*. **(A)** The relative expression of γ1-GLT was analyzed by RT-PCR in sorted populations of Fo B cells, early activated B cells, and GC B cells. Ctrl, black; MIZ1^KO^, red. The expression level of γ1-GLT was normalized to the endogenous Actb level. **(B)** Top, gating strategy for IgG1^neg^IgM^neg^ B GC B cells generated from donor-derived reporter positive B cells. Bottom, graphs displaying cumulative data of percentages for IgG1^neg^IgM^neg^ activated B cells (D4) and GC B cells (D5-7), Ctrl, black; MIZ1^KO^, red. Each symbol (A: Fo Ctrl n = 2, MIZ1^KO^ n = 2; Acti-B Ctrl n = 6, MIZ1^KO^ n = 3; GCB Ctrl = 3, MIZ1^KO^ n = 3; B: day 4 Ctrl n = 8, MIZ1^KO^ n = 8; day 5-6 Ctrl n = 6, MIZ1^KO^ n = 6; day 7 eYFP n = 8, MIZ1^KO^ n = 8;) represents an individual mouse; small horizontal lines show mean and SEM. Data in (A) is from two independent experiments. Data in (B) D4-7 is data from two independent experiments. *, P ≤ 0.05; **, P ≤ 0.01; ***, P ≤ 0.001; ****, P ≤ 0.0001 (multiple t test (B)). ns, not significant.

**Video S2**

Dynamic measurement of Ca^2+^ flux by live cell imaging. Fura-2 stained iGB cells were stimulated with anti-Ig antibody. Ratiometric video images were generated by ImageJ.

**Figure. S4.**
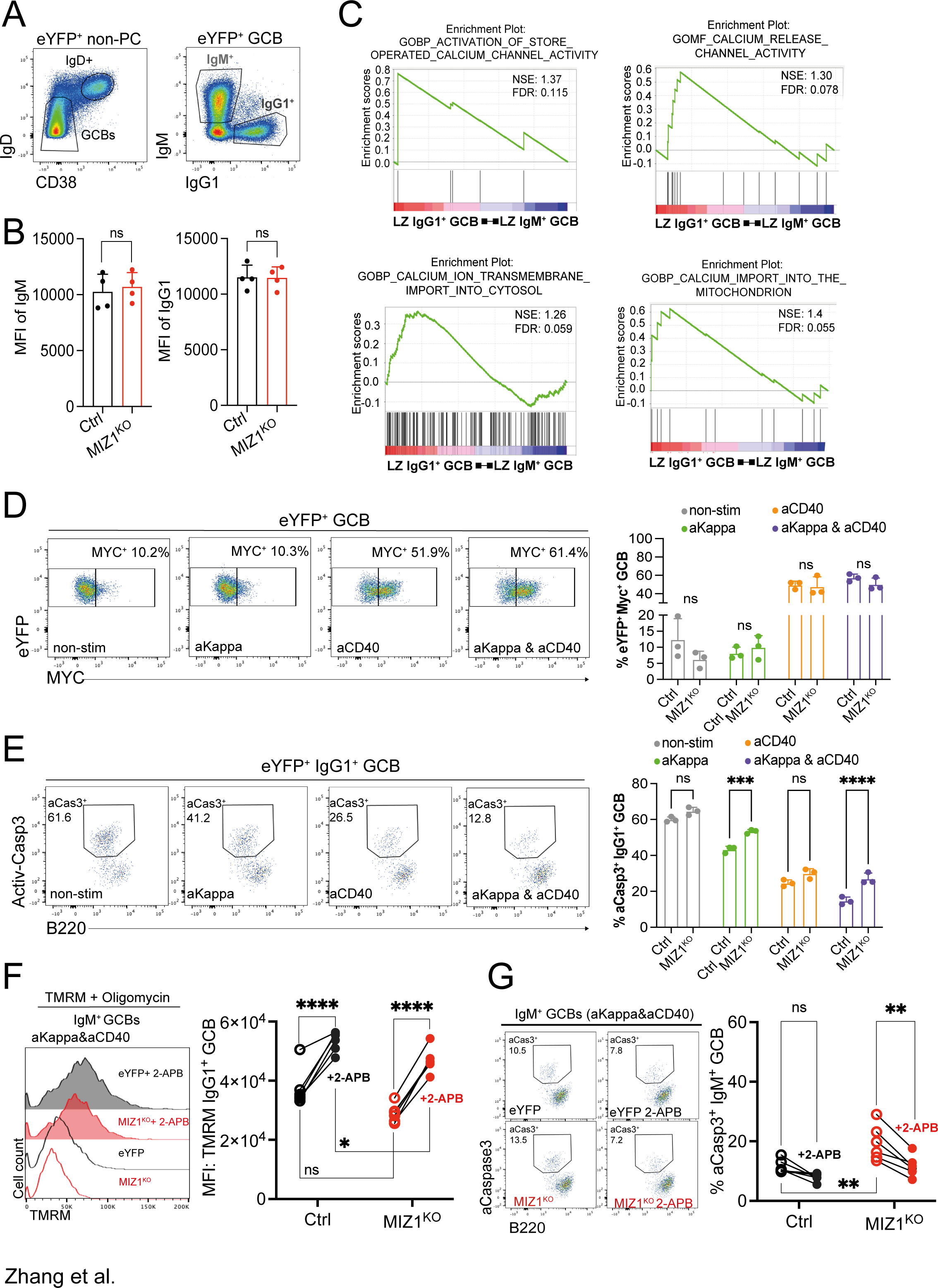
Miz1^KO^ IgG1^+^ GC B cells display irreparable damage despite TFH-help. **(A)** Gating strategy of *ex-vivo* derived IgM^+^ and IgG1^+^ GC B cells subsets, for data presented in (Figure 6 **C-D**). Splenocytes were collected from control and MIZ1^KO^ mice after 8-9 days of 1x 10^9^ SRBCs immunization. Cells were stained with Indo-1 and *ex-vivo* stimulated with anti-Ig antibody (10 ug/ml) during sample acquiring for calcium flux analysis. **(B)** Representative data for BCR expression levels in IgM^+^ and IgG1^+^ GC B cells in (A), measured by gMFI. **(C)** Gene set enrichment analysis (GSEA) of differentially expressed genes (DEGs) in LZ IgG1^+^ versus LZ IgM^+^ GC B cells for signatures associated with store operated Ca^2+^ entry (SOCE), Ca^2+^ release, Ca^2+^ import into cytosol and mitochondrion. FDR, false discovery rate; NES, normalized enrichment score. The list of DEGs was generated from the analysis of RNAseq data in GSE76864. **(D)** Representative plots and percentages of MYC^+^ cells within GC B cells from different *in vitro* stimulation conditions for 3 hours. Non-stimulation condition (non-stim, gray); F(ab’)2 anti-Kappa (aKappa, green), anti-CD40 antibody (aCD40, orange), F(ab’)2 anti-Kappa and anti-CD40 antibody (aKappa & aCD40, purple). **(E)** Representative gating and percentages of active-Caspase3^+^ (aCasp3^+^) cells within IgG1^+^ GC B cells from different *ex vivo* stimulation conditions for 3 hours. Non-stimulation condition (non-stim, gray); F(ab’)2 anti-Kappa (aKappa, green), anti-CD40 antibody (aCD40, orange), F(ab’)2 anti-Kappa and anti-CD40 antibody (aKappa & aCD40, purple). **(F)** Impact of IP3R inhibitor 2-APB on ΔΨ_M_ of *ex vivo* F(ab’)2 anti-Kappa and anti-CD40 antibody stimulated IgM^+^ GC B cells. (G) Impact of IP3R inhibitor 2-APB on apoptosis of *ex vivo* F(ab’)2 anti-Kappa and anti-CD40 antibody stimulated IgM^+^ GC B cells. Each symbol (B: Ctrl n = 4, MIZ1^KO^ n = 4; D: Ctrl n = 3, MIZ1^KO^ n = 3; F: Ctrl n = 6, MIZ1^KO^ n = 6; G: Ctrl n = 6, MIZ1^KO^ n = 6) represents an individual mouse; small horizontal lines show mean and SEM. Data in (B) is representative of three experiments. Data in (D-E) is representative from three independent experiments. Data in (F-G) is from two independent experiments. *, P ≤ 0.05; **, P ≤ 0.01; ***, P ≤ 0.001; ****, P ≤ 0.0001 (unpaired two-tailed Student’s t test (B), Two-way ANOVA (D-G)). ns, not significant.

**Figure. S5.**
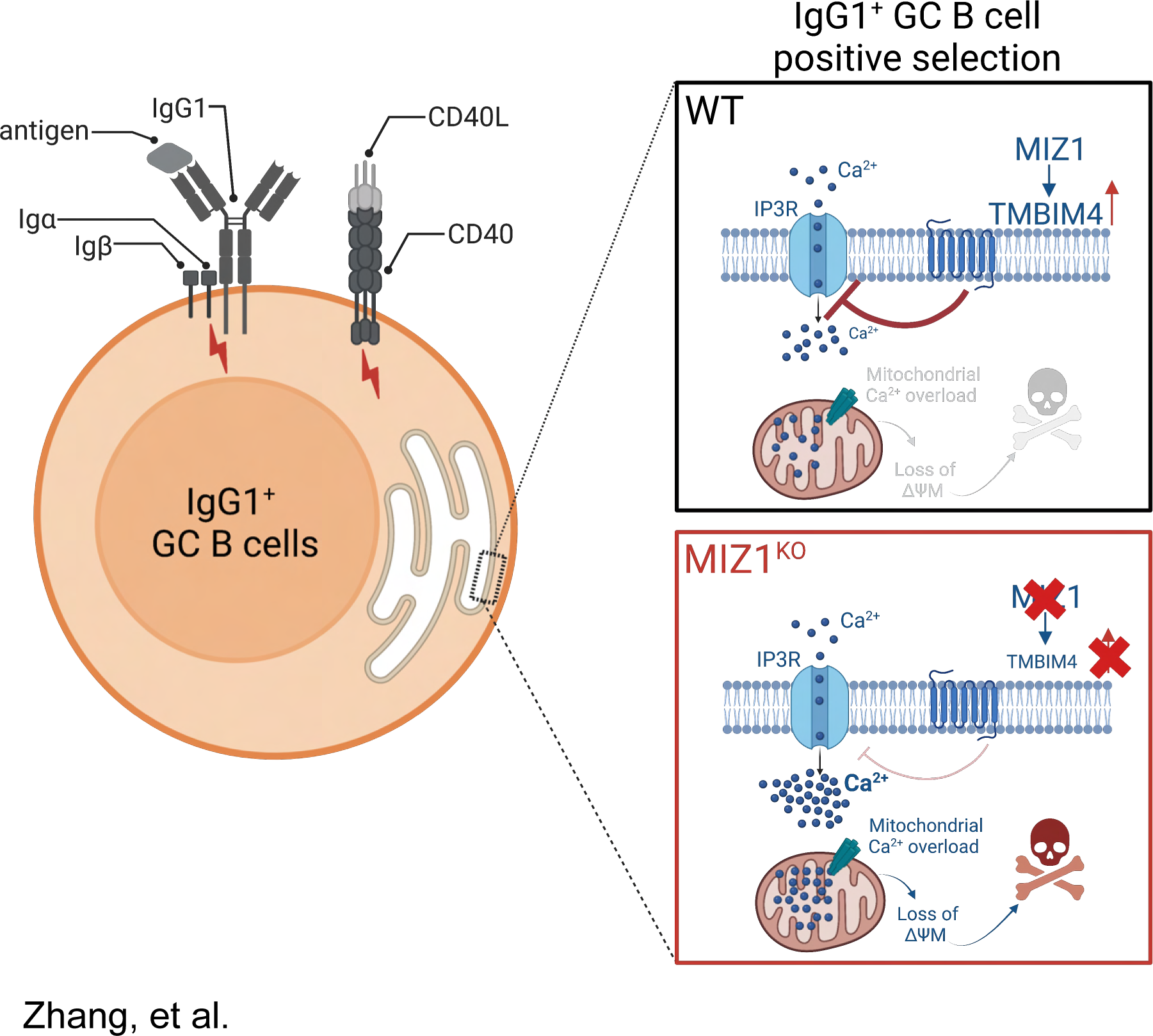
MIZ1-TIMBIM4 safeguards IgG1^+^ GC B cell positive selection. GC B cells undergo a selection process based in part on the affinity of their B cell receptors (BCRs) for antigen immune complexes presented by follicular dendritic cells (FDCs). Positively selected GC B cells (∼5-10% of the LZ) represent those that have productively extracted antigen from FDCs via their BCR and have presented it as peptides on major histocompatibility complex (pMHC) eliciting T follicular helper cell (TFH)-help in the form of CD40 ligand and cytokines. Upon receiving activation signals from TFH, GC B cells highly express MYC and MIZ1. MIZ1 activated the expression of TMBIM4 that effectively regulates Ca^2+^ release from the inositol trisphosphate receptor (IP3R) downstream of IgG1 BCR. Lack of the MIZ1-TMBIM4 axis, led to exacerbated Ca^2+^ mobilization specifically in IgG1^+^ GC B cells, resulting in mitochondrial dysfunction and cell death. Created with BioRender.com.

## Material and methods

### Mice

*Miz1 KO*, *Cγ1-cre*, *R26eYFP^stopFL^* and the transgenic SWHEL allelic BCR system mouse strains have been previously described^71–74^. Mice were maintained on the C57BL/6 background and bred at The Francis Crick Institute biological resources facility under specific pathogen-free conditions. Animal experiments were carried out in accordance with national and institutional guidelines for animal care and were approved by The Francis Crick Institute biological resources facility strategic oversight committee (incorporating the Animal Welfare and Ethical Review Body) and by the Home Office, UK.

### Antibodies and reagents

PE Rabbit Anti-Mouse Caspase 3 (Clone C92-605; BD Biosciences; Cat#550821 RRID: AB_393906); Biotin Rabbit Anti-Mouse Caspase 3 (Clone C92-605; BD Biosciences; Cat#550557 RRID: AB_393750); PerCP/Cyanine5.5 Rat Anti-Mouse CD19 (Clone 6D5; BioLegend; Cat#115534; RRID: AB_2072925); BUV395 Rat Anti-Mouse CD19 (Clone 1D3; BD Biosciences; Cat# 563557; RRID: AB_2722495); Alexa Fluor™ 700 Rat Anti-Mouse CD38 (Clone 90; ThermoFisher(Ebioscience); Cat#56-0381; RRID: AB_657740) BV650 Rat Anti-Mouse CD38 (Clone 90; BD Biosciences; Cat#740489; RRID: AB_2740212) BV650 Rat Anti-Mouse CD45R/B220 (Clone RA3-6B2; BioLegend; Cat#103241; RRID: AB_11204069) BV421™ anti-mouse/human CD45R/B220 Antibody (Clone RA3-6B2; BioLegend; Cat#103251; RRID: AB_2562905) BV605 Mouse Anti-Mouse CD45.1 (Clone A20; BioLegend; Cat#110738; RRID: AB_2562565) BUV395 Mouse Anti-Mouse CD45.2 (Clone 104; BD Biosciences; Cat#564616; RRID: AB_2738867) BV605 Rat Anti-Mouse CD86 (Clone GL1; BioLegend; Cat#105037; RRID: AB_11204429) BV510 Armenian Hamster Anti-Mouse CD95 (Clone Jo2; BD Biosciences; Cat#563646; RRID: AB_2738345) BV421 Armenian Hamster Anti-Mouse CD95 (Clone Jo2; BD Biosciences; Cat#562633; RRID: AB_2737690) BV786 Rat Anti-Mouse CD138 (Syndecan-1) (Clone 281-2; BD Biosciences; Cat#740880; RRID: AB_2740530) Biotin Rat Anti-Mouse CD138 (Syndecan-1) (Clone 281-2; BD Biosciences; Cat#553713; RRID: AB_394999) PerCP-eFluor™ 710 Rat Anti-Mouse CD184 (CXCR4) (Clone 2B11; ThermoFisher(Ebioscience); Cat#46-9991-82; RRID: AB_10670489) PerCP/Cyanine5.5 anti-mouse IgD Antibody (Clone 11-26c.2a; BioLegend; Cat#405710; RRID: AB_1575113) Alexa Fluor® 700 anti-mouse IgD Antibody (Clone 11-26c.2a; BioLegend; Cat#405730; RRID: AB_2563341) Alexa Fluor® 488 anti-mouse IgD Antibody (Clone 11-26c.2a; BioLegend; Cat#405718; RRID: AB_10730619) BV421 anti-mouse IgD Antibody (Clone 11-26c.2a; BioLegend; Cat#405725; RRID: AB_2562743) BUV395 Rat Anti-Mouse IgD (Clone 11-26c.2a; BD Biosciences; Cat#564274; RRID: AB_2738723) BV421 Rat Anti-Mouse CD35 (Clone 8C12; BD Biosciences; Cat#740029; RRID: AB_2739801) Alexa Fluor® 647 Mouse anti-Bcl-6 (Clone K112-91; BD Biosciences; Cat#561525; RRID: AB_10898007) BV421 Rat Anti-Mouse IgG1 (Clone A85-1; BD Biosciences; Cat#562580; RRID: AB_2737664) BUV737 Rat Anti-Mouse IgG1 (Clone A85-1; BD Biosciences; Cat#741733; RRID: AB_2871103) Biotin Rat Anti-Mouse IgG1 (Clone A85-1; BD Biosciences; Cat#550331; RRID: AB_2296342); Fab Fragment Goat Anti-Mouse IgG1 (Jackson ImmunoResearch; Cat#115-477-185; RRID: AB_2632529); Goat Anti-Mouse IgG1, Human ads-AP (Southern Biotech; Cat#1070-04; RRID: AB_2794411) PE-Cyanine7 Rat Anti-Mouse IgM (Clone II/41; ThermoFisher(Ebioscience); Cat#25-5790; RRID: AB_469655) Alexa Fluor® 647 Fab Fragment Goat Anti-Mouse IgM, µ chain specific (Jackson ImmunoResearch; Cat#115-607-020; RRID: AB_2338932) R-Phycoerythrin AffiniPure Fab Fragment Goat Anti-Mouse IgM, µ chain specific (Jackson ImmunoResearch; Cat#115-117-020; RRID: AB_2632508) Goat Anti-Mouse IgM, Human ads-AP (Southern Biotech; Cat#1020-04; RRID: AB_2794200) Mouse Anti-HEL (R. Brink Clone HyHEL9) Alexa Fluor® 647 Anti-c-Myc antibody (Clone Y69; Abcam; Cat#ab190560) Alexa Fluor® 555 Anti-c-Myc antibody (Clone Y69; Abcam; Cat#ab201780) Alexa Fluor® 594 Anti-c-Myc antibody (Clone Y69; Abcam; Cat#ab201775) ZBTB17 Monoclonal Antibody (Clone OTI7G8; Thermo Fisher Scientific; Cat#TA811358) Rabbit anti-TMBIM4 (Thermo Fisher Scientific; Cat#PA5-67464; RRID: AB_2664835) Human Miz-1/ZBTB17 Antibody (R&D; Cat# AF3760) AffiniPure F(ab’)₂ Fragment Goat Anti-Mouse IgG + IgM (H+L) (Jackson ImmunoResearch; Cat#115-006-068; RRID: AB_2338471) Goat Anti-Mouse Ig kappa chain Antibody, F(ab’)2 (Sigma-Aldrich; Cat#AQ500) Armenian Hamster Anti-CD40 (Clone HM40-3; ThermoFisher(Ebioscience); Cat#14-0402-82; RRID: AB_467228); 2-APB (Bio-techne); Recombinant murine IL-4 (Peprotech); Polybrene (Millipore); Poly-L-lysine solution (Sigma-Aldrich); RNeasy Mini Kit (Qiagen); HiSpeed Plasmid Maxi Kit (Qiagen); True-Nuclear Transcription Factor Staining Buffer Set (Biolegend); Fixable Viability Dye (Thermo Fisher Scientific); MitoProbe™ TMRM Kit for Flow Cytometry (Thermo Fisher Scientific); Indo-1, AM (Thermo Fisher Scientific); Fura-2, AM (Thermo Fisher Scientific).

### Immunization, adoptive transfers, and *in vivo* treatments

for immunization in mice with wildtype BCR repertoire, eight-to-12-week-old mice were injected i.v. with 10^9^ defibrinated SRBCs (TCS Bioscience) in PBS (Gibco). 5-ethynyl-2’-deoxyuridine (EdU, Invitrogen) was dissolved in sterile PBS (5 mg/mL); for proliferation studies, 1 mg in a volume of 200 µl was injected i.p. one-hour prior analysis. If not specified, spleens from immunized mice were harvested for analysis after 9 days of immunization. For immunization in SWHEL system, adoptive transfer of HEL^+^ B cells was performed into CD45.1^+^ congenic mice. Briefly, 2×10^5^ HEL-binding B cells together with 2×10^8^ SRBCs conjugated to a specific recombinant HEL protein were injected i.v. into congenic recipients. SRBCs in Alsever’s (TCS Bioscience) were conjugated to recombinant HEL^3X^ (R. Brink) with 1-ethyl-3-(3-dimethylaminopropyl) carbodiimide hydrochloride (Sigma) as previously described^104^. CD45.2^+^ splenic B cells from SW_HEL_ donor mice were purified using anti-mouse CD43 (Ly-48) MicroBeads (Miltenyi Biotec). If not specified, spleens were harvested for analysis after 6 or 7 days after immunization. For FACS analyses and cell FACS sorting, CD45.2^+^ donor splenocytes were enriched by CD45.1 negative selection using anti-mouse biotinylated CD45.1 antibody (clone A20, Ebioscience) and Anti-Biotin MicroBeads (Miltenyi Biotec).

### *Ex vivo* 3D imaging of germinal center B cells

for immunofluorescence staining of GC B cells, GC B cells were enriched by CD43 (clone S7, BD Biosciences), IgD (clone 11-26c, Ebioscience), CD138 (clone 281-2, BD Biosciences), TER-119 (clone TER-119, Biolegend) negative selection using anti-mouse biotinylated antibodies and anti-Biotin Microbeads (Miltenyi Biotec). Freshly isolated GC B cells were washed with PBS and allowed to attach/adhere onto Poly-L-Lysine coated coverslip (Corning, 354085) for 15 min at room temperature (RT) in PBS. Supernatant was gently aspirated, and cells were washed twice with PBS. Cells were fixated with 4% PFA in PBS at RT for 20 min. After fixation, PFA was discard accordingly and cells were washed three times with PBS. Cells were left to dry at room temperature for 30 min before immunostaining. GC B cells were permeabilized and blocked with 0.25% Triton X100 (Sigma-Aldrich) 2% Fetal bovine serum (FBS, Gibco) in PBS for 30 min at RT. Cells were then incubated with Alexa Fluor 647 anti-BCL6 (clone K112-91, BD Biosciences), Alexa Fluor 555 anti-MYC (clone Y69, Abcam) and Alexa Fluor 488 anti-MIZ1 (clone OTI7G8, Thermo Fisher), all diluted at 1:50 in 0.25% Triton X100 2% FBS for 1 hour at RT. Cells were washed three times 5 min each with PBS. Coverslips were mounted on microscope slides (Thermo Fisher Scientific) using SlowFade Glass Antifade Mountant with DAPI (Thermo Fisher Scientific). To perform colocalization analysis spanning whole single cell, z-stack reconstruction of GC B cells was performed using Leica Confocal Microscope SP8 FALCON. Briefly, Navigator from LASX software was used to display the GC B cells on coverslips on field view of 1024×1024 um^2^. Z-step limits up and down were set until DAPI signal was not detectable on focused cells, reaching a z-depth of up to 30um. Individual GC B cells were processed using 3D Crop tool of Imaris (Imaris 9.9.0). Intensity based colocalization of MYC and MIZ1 was performed using ImarisColoc.

### Immunofluorescence on spleen sections

spleens were embedded in optimum cutting temperature compound (Sakura) and flash-frozen in liquid nitrogen. Eight µM to 10 µM tissue sections were cut on an CM3050S cryostat (Leica) and stored at -20 ℃ for future use. On the day of staining, spleen sections were allowed to recover to RT before staining. After this, slides were rehydrated by putting them into a glass tank filled with PBS (Gibco) for 5 min. After this, 4% PFA (Thermo Fisher) was added into each spleen section and incubated at RT for 20 mins to fix the tissue, followed by blocking in PBS with 2% Bovine serum albumin (BSA, Thermo Fisher) and Fc block antibodies (BD Biosciences). For intracellular staining of MYC and MIZ1, 0.25% Triton X-100 (Sigma-Aldrich) in blocking buffer was added to permeabilizing fixed spleen sections at RT for another 20 mins. Intracellular antibodies for MYC (clone Y69, Abcam) and MIZ1 (clone OTI7G8, Thermo Fisher) were diluted in blocking buffer and added onto each section, incubate at 4 ℃, overnight. On second day, wash the slides in PBS for 5 mins, twice. Then apply surface staining antibodies IgD (clone 11-26c, Ebioscience) and CD35 (Clone 8C12, BD Biosciences) onto each spleen sections, incubate at RT for 2 hours with gentle shaking. After staining, slides were washed with PBS (Gibco) and mounted with Antifade Mountant (Invitrogen), covered by coverslip. Images were acquired with Leica SP8 FALCON confocal machine. Image J (v2.3.1) was used to quantify MIZ1^+^ cells with MYC^+^ and MYC^-^ cells within GC area.

### Flow cytometry

single-cell suspensions of spleen were prepared in FACS buffer (2% FBS, 2 mM EDTA, in PBS (Gibco)) and were treated with ACK buffer (Gibco) for erythrocyte lysis. Single-cell suspensions were stained with antibodies. The use of biotinylated antibodies was followed by incubation with fluorochrome labelled streptavidin (1/200 dilution). For the analyses of SW_HEL_ mice, HEL-binding cells were stained with 50 ng/mL HEL (Sigma) or HEL3x (R. Brink) together with anti-mouse IgG1 antibody as the first step, followed by blocking with normal mouse serum (Sigma) before incubating with anti-HEL antibody HyHEL9-A647 (R. Brink). Dead cells were excluded using Zombie NIR™ Fixable Viability Kit (BioLegend). For detection of EdU incorporation, cells were fixed for 15 min at room temperature in 4% paraformaldehyde (Thermo Fisher) after surface marker and viability dye staining. Fixation was followed by Click-iT Plus EdU pacific blue flow cytometry assay kit (Life Technologies) as indicated by supplier. For intracellular staining, samples were fixed for 15 min at room temperature in 4% paraformaldehyde (Thermo Fisher) after surface marker and viability dye staining, followed by intracellular staining with True-Nuclear transcription factor buffer set (Biolegend) for further fixation and permeabilization, as per manufacturer’s specifications. To prevent cross-reactivity, samples were blocked with 10% normal rabbit serum (Thermo Fisher). Samples were acquired on an LSR-Fortessa (BD Biosciences) with FACS-Diva software (BD Biosciences) and data were analyzed with FlowJo software (v10.8.1, Tree Star).

### ELISA detection of HEL-binding serum antibodies

ELISA was performed as described^105^. Briefly, 96-well plates (Corning) were coated with 10 μg/mL HEL in coating buffer and blocked with PBS containing 1% BSA. Serum samples were serially diluted by blocking buffer and were incubated on the plates. After incubation, plates were washed in wash buffer (PBS-0.05%Tween 20) and incubated with AP-conjugated secondary antibodies (anti-IgM, anti-IgG1, Southern Biotech). Plates were washed and followed by adding alkaline phosphatase (AP), p-nitrophenyl phosphate substrate (Sigma-Aldrich). The absorbance was read at 405 nm by using a Microplate Reader (Spectramax 190, Molecular Devices). OD values were exported and were plotted against dilution in smoothed lines. Relative antibody titres were read as maximal dilution where OD was above an arbitrary threshold.

### RT-PCR analysis

total RNA from 10,000 cells was purified using a RNeasy Mini Kit (QIAGEN) according to the manufacturer’s instructions. DNase treatment was performed on-column using RNase-Free DNase Set (QIAGEN). cDNA was generated from 200ng-1ug RNA by using SuperScript™ IV VILO™ Master Mix with ezDNase™ Enzyme Kit (Invitrogen) according to the manufacturer’s instruction. RT-PCR analyses were conducted for target gene γ1-GLT (FAM-labeled probe; Forward primer: 5’-CGAGAAGCCTGAGGAATGTGT-3’ Reverse primer: 5’-GGAGTTAGTTTGGGCAGCAGAT-3’ P: 5’-FAM-TGGTTCTCTCAACCTGTAGTCCATGCCA-3’; ThermoFisher) and ACTB (Actin, Beta) (VIC™/MGB probe, primer limited; Cat#4352341E TaqMan Assay ID: Mm00607939_s1) as reference gene. Samples were measured in duplicates using TaqMan Gene Expression Master Mix (Thermo Fisher) in an Applied Biosystems 7900HT Fast Real-Time machine (Thermo Fisher) with the following thermocycler condition: 50°C for 2 min (1 cycle); 95°C for 10 min (1 cycle); 40 cycles of 95°C for 15 s and 60°C for 1 min. These assays were adopted from a previously published paper^22^. Data is expressed as a fold-change using the ΔΔC_T_ method.

### *In vitro* GC system

the *in vitro* GC (iGC) system was slightly modified from previously published protocol^106^. 40LB feeder cells for iGC culture were obtained from the lab of Daisuke Kitamura. These cells were maintained by G418 (Gibco) and puromycin (Gibco) selection. Briefly, one day before cell seeding, 0.5 x 10^6^ of 40LB cells were plated in each well of one 6-well plate and allowed to attach overnight. Before seeding B cells, feeder cells were treated with Mitomycin C (Apexbio Technology) at the concentration of 10 μg/mL for 2 hours and then washed with PBS. After this, 0.17 x 10^6^ purified naïve B cells were plated in each well of a 6-well plate on top of 40LB feeder cells in 6 mL of B cell culture medium. (DMEM media (Sigma) supplemented with 10% heat-inactivated fetal bovine serum (FBS) (Thermo Fisher), 100μM non-essential amino acids (Thermo Fisher), 2mM L-Glutamine (Thermo Fisher), 50μM 2-Mercaptoethanol (Thermo Fisher) and Penicillin-Streptomycin (GE Healthcare Life Sciences).) supplemented with 1ng/mL recombinant mouse IL-4 (Peprotech)). Cells were incubated at 37°C, 5% CO_2_ for 4-7 days for the generation of induced GC B cells.

### *Ex vivo* stimulation of GC B cells

mice were i.v. immunized with 1 x 10^9^ SRBCs and their spleens were collected after 8-9 days of immunization. RBCs depleted splenocytes from these mice were *ex vivo* stimulated with F(ab’)2 anti-Kappa (1 ug/ml, Sigma-Aldrich) and (or) anti-CD40 (0.5 ug/ml, Thermo Fisher) antibodies for 3 hours and stained for TMRM (Thermo Fisher) and active-Caspase3 (BD Biosciences). To directly test the impact of calcium flux on GC B cell death, IP3R inhibitor 2-APB was introduced into the *ex vivo* culture system. Cells were firstly *ex vivo* treated with 2-APB (100 μM, Bio-techne) for 30 mins and followed by stimulation with 100uM. F(ab’)2 anti-Kappa (1 μg/ml) and anti-CD40 (0.5 μg/ml) antibodies. Splenocytes were stimulated in the presence of 2-APB for another 3 hours and collected to stain for TMRM or active-Caspase3 by flow cytometry.

### Calcium flux

To analyze calcium flux by flow cytometry, spleens were taken from SRBCs immunized mice, 8-9 days after immunization. Spleens were smashed on 70 um filter and single cell suspensions from spleens were spun down and supernatants were discarded. 2 ml ACK buffer (Sigma-Aldrich) was used to lyse red blood cells (RBCs). 1 min after lysis, the same volume of complete culture medium was added to stop the reaction and spun down again (repeated this step until RBCs have been lysed completely). After this, cells were spun down and re-suspended with HBSS buffer at the concentration of 2 x 10^6^ cells / ml. DMSO dissolved Indo-1 (Invitrogen, 1mM stock) was added to the cell suspension to a final concentration of 1μM / ml. Cell suspensions were left in the dark at RT for 30 min to load Indo-1. The reaction was stopped by adding twice the volume of HBSS-10% FBS. Cell suspensions were spun down and re-suspended with 200 μl HBSS-10% FBS again to allow complete de-esterification of extra AM moieties, Indo-1or Fura-2 (37 °C, 5% CO_2_ for 30 min). Cells were then labelled with anti-CD43 antibody as a dump marker in FACS buffer at 4 °C for 30 min, washed once and were “calmed” by culturing in complete culture medium at 37 °C, 5% CO_2_ for 1-2 hours (Khalil, Cambier et al. 2012). After calming, cells were processed and detected by flow cytometry. Stimulation through BCRs were done by adding 10 ug/ml F(ab’)^2^ anti-Kappa (Sigma-Aldrich) or F(ab’)^2^ anti-mouse IgG+IgM (H+L) (Jackson ImmunoResearch) during sample acquisition. Samples were analyzed by BD LSRFortessa Analyzer (BD Biosciences) with BD FACSDiva. Laser line used was 350 nm (UltraViolet), filters were set for 450/50 (calcium bound) and 525/50 (calcium free). Raw data was exported and the ratio of bound to free calcium was calculated in FlowJo (FlowJo, LLC) as a Derived Parameter.

To analyze calcium flux by live imaging, iGB cells were collected after 6-7 days on iGC culture and were stained with Fura-2 (Invitrogen) at 1μM / ml. 4-Chamber Glass Bottom Dish (Cellvis) was pre-treated with Poly-L-lysine solution (Sigma). Before seeding iGB cells, Poly-L-lysine solution was discarded, and the dishes were left to dry at RT for 15 mins. 200 μl Fura-2 stained iGB cells in HBSS containing10% FBS were seeded into each well at the concentration of 5-10 x 10^6^ cells /ml. Cells were left at RT for at least 10 mins to allow them attach to the bottom, and bottom attached iGB cells were imaged by Nikon ECLIPSE Ti2, laser line used was 340 nm (calcium bound) and 385 (calcium free). Ratiomatric images (ratio: bound / free) were processed and generated by ImageJ (v. 2.9.0).

### Gene expression analysis, chromatin immunoprecipitation, and sequencing

For the gene expression profiling of eYFP and MIZ1^KO^ LZ GC B cells (CD138^neg^, B220^+^, CD19^+^ CD38^low^, Fas^high^, Cxcr4^low^ CD86^high^) were sorted by flow cytometry at day 10 after immunization with SRBCs using a FACSAria III or a FACSAria Fusion (BD Biosciences). Cells were sorted in RTL Buffer Plus (Qiagen) containing 1% ß-mercaptoethanol (Sigma) and RNA purified using AllPrep DNA/RNA Mini and Micro Kits (Qiagen) following manufacturer’s instructions. RNA-sequencing was performed at The Francis Crick Institute advanced sequencing facility.

The RSEM package (v1.3.0)^107^ and STAR (v2.5.2a)^108^ were used to align reads to the mouse mm10 transcriptome, taken from Ensembl (v. GRCm38) and available at UCSC (https://genome.ucsc.edu). For RSEM, all parameters were run as default using the “–forward-prob 0” option for a strand specific protocol. Differential expression analysis was carried out with DESeq2 package (v.1.18.1)^109^ within R (v3.4.3)^110^ (http://www.R-project.org). Genes were differentially expressed if adjusted p-value (AdjP) was less than 0.05. All the analyses for RNA-seq-generated expression profiles were done with ranked gene lists using the Wald statistics.

For chromatin immunoprecipitation, Raji and P493-6 cells were cultured at 37°C (5% CO_2_) in DMEM supplemented GlutaMAX™, non-essential aminoacids, penicillin-streptomycin, HEPES (Gibco), ß-mercaptoethanol (Sigma) and 10% FBS (Thermo Fisher)). For P493-6 cells, doxycycline and β-oestradiol was added into culture medium to induce the expression of MYC. ChIP protocol was adapted from^111^. Briefly, cells in B cell media were fixed with formaldehyde (Sigma) at final concentration of 1% for 10 minutes at room temperature. Glycine (Fisher Scientific) was added at a final concentration of 0.125M during 5 minutes at room temperature. Cells were washed, pelleted, and stored at -80 °C. After cell lysis, the DNA of 50 x10^6^ cells was fragmented using BIORUPTOR 200 immersion sonicator under the following conditions: high Power, 30 sec ON, 30 sec OFF (40 Cycles or less). 100uL of Pierce Protein A/G Magnetic Beads were coupled to 10 µg of antibody or control rabbit IgG antibody (Sigma). DNA was incubated with antibody-coupled beads and after uncoupling, DNA was purified with NucleoSpin Gel and PCR Clean-up (Macheray-Nagel) following manufacturer’s protocol for SDS-rich samples, using Buffer NTB (Macheray-Nagel). Samples were sequenced on an Illumina HiSeq2500 generating 100bp single ended reads. ChIP-seq reads were aligned to the mouse mm10 genome assembly using BWA version 0.7.15^112^ with a maximum mismatch of 2 bases. Picard tools version 2.1.1 (www.broadinstitute.github.io/picard) was used to sort, mark duplicates and index the resulting alignment bam files. Files were normalized and “tdf” files for visualization purposes were created using IGVtools software version 2.3.75 (http://www.broadinstitute.org/igv) by extending reads by 50 bp and normalizing to 10 million mapped reads per sample. Peaks were called using the standard parameters by comparing immunoprecipitated samples to their respective input and/or IgG controls using MACS version 2.1.1^113^. Peaks called by MACS were annotated using the ‘annotatepeaks’ function in the Homer version 4.8.3 software package^114^. Common and unique peaks across experiments were determined using a custom script.

### Retroviral constructs and transductions

Murine *Tmbim4* expression construct was made by inserting open reading frame into the MSCV retroviral vector followed by an 2A self-cleaving peptides T2A (thosea asigna virus 2A) and *RFP* (or *Tagblue*) as an expression marker. All retroviral vectors were generated by Vectorbuilder. Retroviral transduction was modified from the published protocol (Chapter 13, Rinako et. al., Germinal center methods and protocols, Springer Protocols). Retrovirus was generated by transfecting the Plat-E packaging cell line with 10 µg plasmid DNA and 30 µl FuGene HD transfection reagent (Promega). Viral supernatants were collected from transfected Plat-E cells at 40, 64, 88 and 112 hours after transfection and mixed with 1/4 of original supernatant volume of PEG-it viral preparation solution (SBI Systems Biosciences). Mixture was stored at 4 °C until the day of infection and was centrifuged at 1500 g for 30 min at 4 °C to concentrate retrovirus. 200 µl HEL (20 mg/ml, Sigma-Aldrich) was injected i.p. into donor mice. Six hours after injection, spleens from donor mice were collected and B cells enriched by negative selection. B cells were stimulated in vitro with anti-CD40 antibody (clone HM40-3, Biolegend) overnight to fully activate HEL^+^ B cells. On the second day, B cells were spin-infected with concentrated retrovirus at 750 g for 90 min at 30 °C. Transduced donor B cells were then adoptively transferred into CD45.1^+^ congenic recipient mice.

### Quantification and statistical analysis

Data were analyzed with unpaired two-tailed Student’s t test; a p-value of 0.05 or less was considered significant. Prism (v9.4.1, GraphPad) was used for statistical analysis. A single asterisk (^∗^) in the graphs of figures represents a p-value ≤0.05, double asterisks (^∗∗^) a p-value ≤0.01, triple asterisks (^∗∗∗^) a p-value ≤0.001, quadruple asterisks (^∗∗∗∗^) a p-value ≤0.0001, and “ns” stands for non-statistically significant, i.e., a p-value >0.05. Data in text and figures are represented as median or mean ± standard error of the mean (SEM), each case is indicated in figure legends.

### Online supplemental material

RNAseq data of LZ_Miz1ff; LZ_eYFP (This paper; GSE234361; access token: spwdakyajfmnzyz); CHIP-seq data of MIZ1 binding sites in P493-6 and Raji (This paper; GSE234360; access token: yzcdcumkrdwblkj); CHIP-seq data of MIZ1 binding sites in primary murine B cells((de Pretis S, et al., 2017); GSE98419); Gene set enrichment analysis (GSEA) of differentially expressed genes (DEGs) in LZ IgG1+ versus LZ IgM+ GC B cells ((Gitlin AD, et al., 2016); GSE76864)

## Notes

### Competing Interest Statement

The authors have declared no competing interest.

